# Aberrant E-I Balance and Brain Criticality in Major Depressive Disorder

**DOI:** 10.1101/2025.10.19.683280

**Authors:** Asako Mitsuto, Abhimanyu Bhardwaj, Linden Parkes, Fatima Faruqi, Linda Carpenter, Andrew Westbrook

## Abstract

Brain criticality and complexity are increasingly recognized as promising biomarkers for psychiatric disorders. In Major Depressive Disorder (MDD), disordered neural dynamics have been reported, but their nature and consistency remain incompletely understood. Here, we study brain criticality, excitation–inhibition (E/I) balance, combined excitation– inhibition strength (E+I), and complexity of brain dynamics associated with MDD. Using resting-state EEG from 183 patients with MDD and 133 healthy controls (HC), we identified disruptions of critical dynamics and excitation–inhibition balance which discriminate groups. We found that amplitude bistability is lower, and long-range temporal correlations are weaker in MDD, implying deviation from criticality. Excitation-inhibition metrics show frequency-specific alterations in MDD. Estimates of excitation-inhibition ratios (E/I) derived from the statistical properties of amplitude fluctuations show higher values in HC than MDD in the θ band, indicating relative over-excitation, and lower values in the γ band, indicating relative over-inhibition. An excitation– inhibition strength index reflecting combined excitatory and inhibitory drive (E+I), was decreased in θ through β bands and increased in γ in MDD. Collectively, excitation-inhibition measures suggest decreased inhibitory drive in the mechanisms underlying θ oscillations in MDD and increased inhibitory drive in the mechanisms underlying γ oscillations. Classification using least absolute shrinkage and selection operator (LASSO) regression achieved high accuracy and the predictive feature set includes measures of criticality, E/I ratios, and combined E+I strength. These findings elucidate pathological alterations of brain dynamics in MDD and define a complex system fingerprint, supporting the development of biomarkers for diagnosis and treatment.

## Introduction

Brain criticality, excitation-inhibition (E/I) balance, and complexity have emerged as promising biomarkers for psychiatric disorders, including major depressive disorder (MDD). A central hypothesis is that healthy neuronal networks operate near a critical point, where excitation and inhibition are balanced. Operating near criticality maximizes computational properties, including dynamic range, information storage and transmission, flexibility (Beggs and Plenz 2003, Shew and Plenz 2013, Cocchi, Gollo et al. 2017, Hengen and Shew 2025). These benefits arise from emergent dynamical properties of neuronal populations across spatial and temporal scales maximized near criticality including complexity (or irregularity), susceptibility (responsiveness), and long-range temporal correlations (LRTC; persistence) (Linkenkaer-Hansen, Nikouline et al. 2001, Buzsáki and Draguhn 2004, Kim, Lee et al. 2020, Kim and Lee 2020, Lotfi, Feliciano et al. 2021). Deviations from criticality are theoretically and experimentally linked to impaired information processing. Clinical symptoms across a diversity of disorders may reflect divergence from criticality, including the cognitive deficits that characterize patients with MDD (Zimmern 2020, O’Byrne and Jerbi 2022, Hengen and Shew 2025).

Prior studies of E/I balance and criticality in MDD report conflicting findings. In some studies, MDD brains deviate from criticality towards either excitation- or inhibition-dominance (Linkenkaer-Hansen, Monto et al. 2005, Hengen and Shew 2025, Momi, Wang et al. 2025). Other studies report stronger emergent properties, including stronger long-range temporal correlations (LRTC) in comparison with healthy controls (HC) (Lee, Yang et al. 2007, Gärtner, Irrmischer et al. 2017) implying that MDD brains operate closer to criticality. Conflicting results may reflect real distinctions in mechanism-specific E/I ratios in MDD versus HC. Indeed, postmortem and magnetic resonance spectroscopy (MRS) studies demonstrate widespread markers of excitation and inhibition that differ by brain region, in MDD versus HC. MDD patients, for example, have decreased GABAergic and increased glutamatergic signaling markers, corresponding to increased excitation in the default network and increased inhibition in the lateral prefrontal cortex (Dubin, Mao et al. 2016, Godfrey, Gardner et al. 2018, Hu, Tan et al. 2023). Moreover, individual differences in depression symptom severity, and the antidepressant treatment efficacy in non-invasive brain stimulation studies correlate with mechanism-specific indices of E/I balance and brain criticality inferred from dynamical properties (Radhu et al. 2013, Xin et al., 2022, Dhami et al, 2023, Bhardwaj et al, 2023). These findings suggest that complementary, mechanism-specific indices of E/I balance and criticality are useful for distinguishing MDD patients. An alternative explanation for conflicting results across prior studies may be low reliability: most prior studies contrasting MDD patients and HC involved small sample sizes (e.g. fewer than 30 participants). To address conflicting results and establish robust, generalizable, neurophysiological biomarkers of MDD, it is crucial to investigate how criticality and E/I balance jointly shape brain dynamics through large-sample studies and convergent analytical methods (Avramiea, Diachenko et al. 2025).

We analyzed a dataset comprising 183 patients with MDD who completed pre-treatment EEG recordings. For comparison, we included 133 HC from the publicly available Max Planck Institute Leipzig Mind-Brain-Body dataset (MPI-LEMON; Babayan et al., 2019), which provides high-quality resting-state EEG, selecting a subset to match the mean age and sex ratio of our MDD group. We employed a multidimensional analysis integrating EEG-derived metrics that capture three distinct aspects of neural dynamics: criticality, excitation–inhibition balance (E/I), and complexity (Comsa 2019, Avramiea, Diachenko et al. 2025).

Criticality was indexed by the bistability index (BiS), which quantifies alternations between high- and low-power states (Freyer, Aquino et al. 2009), and by detrended fluctuation analysis (DFA) (Hardstone, Poil et al. 2012), which estimates long-range temporal correlations. Regarding bistability, Freyer et al. showed that bistable states characterized by a mix of high- and low-amplitude α oscillations emerge near a subcritical Hopf bifurcation, a hallmark of critical transitions (Freyer, Aquino et al. 2009, Freyer, Roberts et al. 2012). This mechanism is generalizable, since metastability based on Hopf bifurcations could apply to other frequency bands as well. DFA captures scale-free long-range temporal correlations, a classic signature of systems poised at criticality (Hardstone, Poil et al. 2012).

E/I balance was assessed using the high-to-low power ratio (E/I_HLP_) (Avramiea, Diachenko et al. 2025), and the functional E/I ratio (fE/I) (Bruining, Hardstone et al. 2020) derived from amplitude–fluctuation coupling. The high-to-low power ratio (E/I_HLP_) is derived from a bi-exponential characterization of amplitude distributions that reflects underlying excitatory–inhibitory (E/I) dynamics in cortical networks (Avramiea, Diachenko et al. 2025). This biomarker has been empirically validated using stereotactic EEG (sEEG) in patients with epilepsy. During ictal periods, E/I_HLP_ was significantly higher in seizure onset zones (SOZ) compared to non-SOZ regions, consistent with localized shifts toward excitation-dominated dynamics. These findings support the use of E/I_HLP_ as a non-invasive, physiologically grounded marker of cortical E/I balance. The functional E/I ratio (fE/I) captures excitation–inhibition balance by quantifying the covariance of oscillation amplitude and short-timescale fluctuations and was shown to decrease relative to placebo after pharmacological enhancement of GABAergic inhibition with zolpidem in healthy adults (Bruining, Hardstone et al. 2020).

Finally, the separation of high- and low-power oscillations (E+I_HLS_) (Avramiea, Diachenko et al. 2025) has been analyzed as an index of combined excitatory and inhibitory drive. E+I_HLS_ has been shown, for example, to differentiate SOZ from non-SOZ electrodes during seizures in sEEG data, particularly in the δ–α bands (Avramiea, Diachenko et al. 2025).

Towards mechanism-specificity, we computed all these measures in logarithmically-spaced frequency-specific bins spanning δ through low γ. Complexity was quantified with Lempel–Ziv complexity (LZC), which captures the irregularity of temporal sequences. Together, these metrics provided a rich characterization of complex brain dynamics, enabling us to test neurophysiological signatures of MDD across domains and to evaluate their potential as biomarkers.

Based on the theoretical considerations and prior empirical evidence, we hypothesized that the MDD brain deviates from criticality; we made no specific assumptions about whether divergence would reflect over-excitation or over-inhibition, relative to controls (Figure 1). We tested whether there is evidence of divergence from criticality and whether it corresponded with a subcritical or supercritical shift across distinct frequency bands. Our hypothesis predicts that criticality indices (BiS and DFA) are reduced in MDD, suggesting divergence from criticality, relative to healthy controls. We also predicted that aberrant criticality would be reflected by shifts in E/I ratios across frequency bands, captured by E/I balance indices (E/I_HLP_ and fE/I), and furthermore that we would gain additional inferential leverage about MDD-related alterations in excitation versus inhibition by evaluating combined excitatory and inhibitory drive (E+I_HLS_). We expected that signal complexity would be reduced in MDD. Because complexity is theoretically coupled to criticality (Toker, Pappas et al. 2022), we predicted that brains with MDD would show lower complexity, serving as a complementary marker of altered critical dynamics.

**Figure 1.**
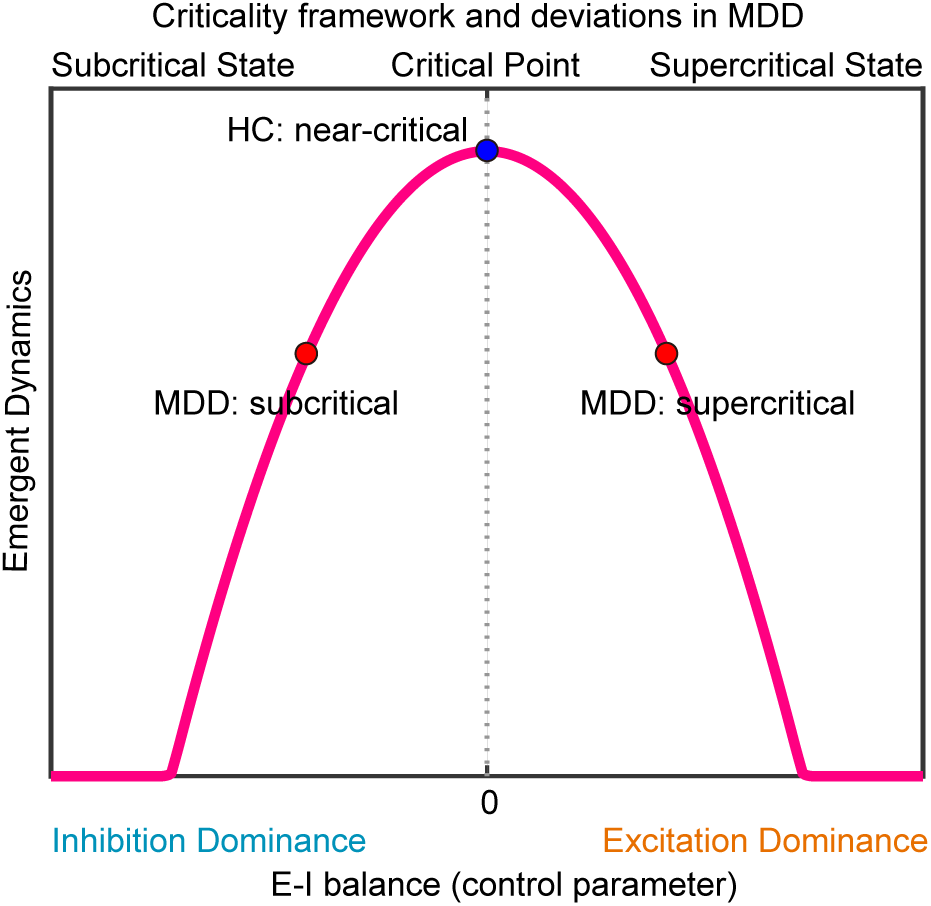
Brain criticality hypothesis in major depressive disorder. Schematic illustration showing excitation–inhibition (E/I) balance as a control parameter for brain criticality. The inverted-U curve depicts emergent dynamics (e.g., complexity, bistability, and long-range temporal correlations), peaking at the critical point where excitation and inhibition are balanced. The blue marker (HC) reflects operation near criticality hypothesized for healthy adult brains. Red markers illustrate hypothesized MDD deviations toward subcritical (inhibition-dominant, left) and supercritical (excitation-dominant, right) regimes. Top labels indicate subcritical, critical, and supercritical states; the x-axis annotates inhibition dominance (−) and excitation dominance (+).

## Material and Methods

### Patients

We analyzed resting-state EEG from 183 patients with MDD presenting for treatment and recruited for a research EEG study at the Butler Hospital TMS Clinic. The full clinical dataset includes 190 unique patients. We selected the EEG recorded at the first pre-treatment session of each patient’s earliest treatment series. Following artifact cleaning, we retained only those patients with at least 4.65 minutes of pre-treatment resting-state EEG, resulting in a final sample of 183.

The final MDD sample had a mean age of 46.8 ± 16.2 years and included 60 males and 123 females (male/female ratio = 0.33). Patients had a primary diagnosis of (nonpsychotic) MDD and of moderate to severe intensity MDD symptoms. Depression symptom severity, assessed with the Inventory of Depressive Symptomatology– Self Report (IDS-SR), averaged 46.7 ± 10.3. The clinical population was considered treatment-resistant, with most patients having failed multiple adequate antidepressant trials and many experiencing chronic, disabling episodes. Nearly all patients were taking stable regimens of psychotropic medications. All participants provided written informed consent, and study procedures were approved by the Butler Hospital Institutional Review Board. Demographic details and IDS-SR scores are summarized in Supplementary Table 1.

### Healthy Controls

We included a comparison sample of HCs drawn from the publicly available Max Planck Institute Leipzig Mind-Brain-Body dataset (MPI-LEMON; Babayan et al., 2019). The LEMON dataset comprises 228 adults recruited in Leipzig, Germany, between 2013 and 2015, with a younger cohort (N=154, 20–35 years) and an older cohort (N=74, 59–77 years). From this resource, we selected 133 participants to approximate the MDD sample in mean age and sex distribution by combining all females, and a subset of male participants. Participants in the HC group were aged 44.9 ± 21.4 years (estimated from 5-year age bins based on midpoint values) and comprised 55 male and 78 female participants. The severity of depression was minimal in the HC sample, with Hamilton Depression Rating Scale scores averaging 2.54 ± 2.61. All controls were free of psychiatric or neurological illness as determined by structured clinical interviews and self-report questionnaires administered in the original study.

### EEG acquisition

Resting-state EEG was acquired from MDD patients (64-channel ANT Neuro, Butler Hospital) and HCs (62-channel ActiCAP, LEMON dataset). To harmonize length and epoch structure, we built a 4.65-min eyes-closed series per participant. For HC, five 1-min eyes-closed epochs were concatenated, then 10.6 second were removed from both the beginning and the end. For MDD, a single ∼5-min eyes-closed recording was split into five 1-min epochs, reordered (1–3–5–2–4), and the same 10.6 second trimming was applied at the start and end. This yielded equal-length, epoch-matched time series across groups for all analyses.

### EEG Preprocessing

EEG preprocessing was performed in EEGLAB (version 2022.0). The EEG data were first high-pass filtered at 1 Hz and down-sampled to 512 Hz. Channels with abnormal activity were identified using the clean_artifacts function (ChannelCriterion = 0.8, LineNoiseCriterion = 4) and removed and interpolated. Data were re-referenced to the average. To attenuate transient artifacts, artifact subspace reconstruction (ASR; threshold = 20) was applied to reconstruct contaminated data segments rather than removing them.

For artifact removal, independent component analysis (ICA) was applied to each cleaned dataset. Components corresponding to muscle, ocular, cardiac, line noise, or channel noise activity were automatically classified with the ICLabel algorithm, and those with >70% artifact label probability were removed. The resulting cleaned data were re-referenced to the average.

### Frequency bands

We examined 13 sub-bands spanning 1–73 Hz, obtained by logarithmically-spaced bins for BiS, DFA, E/I_HLP_, fE/I, and E+I_HLS_.

### Metrics

#### Criticality metrics

##### Bistability (BiS)

The bistability index (BiS) quantifies how strongly neural activity alternates between high- and low-power states, a hallmark of systems operating near criticality (Freyer, Aquino et al. 2009, Freyer, Roberts et al. 2012). In inhibition-dominant or excitation-dominant regimes, oscillations fluctuate smoothly around a single low-power or high-power mode, respectively. In contrast, near the critical regime, the brain exhibits spontaneous switching between low- and high-power modes, resulting in a bimodal distribution of power. BiS quantifies this degree of bimodality, with higher values indicating more clear alternation between these two states and thus stronger signatures of criticality. Detailed methods for computing BiS are described in Supplementary Information.

### Detrended Fluctuation Analysis (DFA)

We estimated long-range temporal correlations from band-limited EEG amplitude envelopes using detrended fluctuation analysis (DFA). For each frequency bin, the envelope was integrated to form a signal profile, which was segmented into windows of length s, advanced with 50% overlap (step = s/2). Within each window, local linear trends were removed, and root-mean-square fluctuations were computed. The DFA exponent (an estimator of the Hurst exponent) was obtained as the slope of the log–log regression of fluctuation versus window size over the fitting interval [*S_min_*, 30 s], where *S_min_* was adjusted per band (Hardstone, Poil et al. 2012).

A practical guideline for DFA is to include at least 6–10 independent windows of the longest time scale (Hardstone, Poil et al. 2012, Diachenko, Sharma et al. 2024). With our sequential minimum 49-second recordings, this criterion is only partially met. However, the full EEG segments (49, 60, 60, 60, and 49 seconds) provide 8 independent windows when using the 30-second window length applied in our analysis.

### Excitation–inhibition balance metrics

#### The proportion of high- and low-power oscillations (E/I_HLP_)

E/I_HLP_ provides an estimate of the excitation–inhibition (E/I) ratio from EEG amplitudes. Greater inhibition produces small-amplitude oscillations, whereas greater excitation produces large-amplitude oscillations. When the system is near criticality, neural dynamics fluctuate between these excitation-dominant and inhibition-dominant states. E/I_HLP_ captures this balance by quantifying the relative predominance of high- versus low-power oscillatory states, offering an electrophysiological proxy of the underlying E/I balance. Details of its computation are described in the Supplementary Information.

### Functional E/I ratio (fE/I)

The functional excitation–inhibition ratio (fE/I) provides a model-based estimate of cortical excitation–inhibition balance by quantifying the correlation between oscillatory amplitude and long-range temporal correlations (LRTC) (Bruining, Hardstone et al. 2020). For each channel × frequency bin, the amplitude envelope was extracted using the Hilbert transform, integrated into a signal profile, and segmented into overlapping 5-second windows. Within each window, fluctuations were normalized by the local mean amplitude, detrended, and quantified as root-mean-square deviations. The fE/I ratio was then defined as one minus the correlation between windowed mean amplitudes and corresponding normalized fluctuations, such that values near 1.0 indicate balanced excitation–inhibition dynamics, whereas deviations above and below 1.0 reflect shifts toward excitation- or inhibition-dominated regimes, respectively. Estimates were only retained when the corresponding DFA exponent exceeded 0.6.

For fE/I, we used fixed 5-second windows with 80% overlap.

### Excitation-inhibition strength

#### Separation of High- and Low- Power Oscillations (E+I_HLS_)

E+I_HLS_ indexes the strength of both excitatory and inhibitory processes. When excitation and inhibition are strong but balanced, neural activity alternates between distinct high- and low-amplitude states, creating a clear separation in the power distribution between them. In contrast, when these forces are weak or imbalanced, the distinction between high- and low-power states becomes less pronounced, and one state may dominate or blur into the other. HLS quantifies this separation, with larger values indicating greater distinction between excitatory and inhibitory states. Details of the computation are provided in the Supplementary Information.

### Complexity metrics

#### Lempel–Ziv complexity (LZC)

LZC was used to quantify the irregularity of EEG signals by counting the number of distinct patterns in the binarized time series (Comsa 2019). Higher normalized LZC indicates greater signal irregularity.

We used the 2024.08.18 release of a package for BiS, DFA, E/I_HLP_, fE/I, and E+I_HLS_ (Hardstone, Poil et al. 2012, Bruining, Hardstone et al. 2020, Avramiea, Diachenko et al. 2025) and open code for LZC (Comsa 2019). All analyses were run with under Python 3.11 on a high-performance computing cluster.

### Statistical analysis for BiS, DFA, fE/I, E/I_HLP_, and E+I_HLS_

Group-level statistical comparisons between MDD and HC were performed separately for each metric × frequency band. For each subject, metric values were averaged across channels. Unadjusted band-wise contrasts were tested with Welch’s unequal-variance t-tests (appropriate with unequal sample sizes and without assuming variance homogeneity). Resulting p-values were adjusted for multiple comparisons across the 13 frequency bands within each metric using the Benjamini–Hochberg false discovery rate (FDR) procedure as implemented in MATLAB’s mafdr function.

To quantify the magnitude of group differences, we calculated group mean differences together with 95% confidence intervals estimated using Welch’s unequal-variance standard error and Satterthwaite’s approximation for the effective degrees of freedom, ensuring valid inference under heterogeneity of variance. In addition, we reported Cohen’s d, defined as the mean difference divided by a pooled standard deviation across groups, with confidence intervals derived from the Hedges–Olkin variance approximation.

To control for residual demographic differences between groups, we performed covariate-adjusted analyses using robust linear regression models. For each band and metric, we fit a robust regression with group (MDD vs. HC) as the predictor of interest, including age (z-scored) and sex (coded male=1, female=0) as covariates. Robustness was achieved using iteratively reweighted least squares, implemented in MATLAB’s fitlm function with RobustOpts enabled. To obtain covariate-adjusted group differences, we extracted group coefficients and their associated t-statistics. In addition, permutation tests were applied in the Freedman–Lane framework, where residuals from reduced models (age + sex only) were permuted and added back to predicted values to generate empirical null distributions of group effects.

These statistical analyses were performed in MATLAB R2023a on Windows 11. Figures displayed band-wise adjusted means ± standard errors, with uncorrected and FDR-corrected significance markers indicated.

### Statistical analysis of LZC

We first extracted per-subject LZC values by averaging across EEG channels. To compare groups, we implemented three complementary approaches.

Unadjusted group comparisons: Mean LZC values were contrasted between MDD and HC using Welch’s t-tests, Mann–Whitney U tests, and permutation tests (10,000 label-swapping iterations). We reported effect sizes as Cohen’s d and visualized distributions with boxplots and histograms.

Covariate-adjusted analyses: To control for demographic confounds, we next fit robust linear regression models (iteratively reweighted least squares, MATLAB fitlm (with RobustOpts=on), with LZC as the dependent variable, group (MDD vs. HC) as the predictor of interest, and age (z-scored) and sex (coded male=1, female=0) as covariates.

Permutation with covariates (Freedman–Lane): Group effects in the covariate-adjusted models were validated via residual-based permutation tests (10,000 permutations). Specifically, residuals from the reduced model (covariates only) were permuted and recombined with fitted values, and the group coefficient t-statistic was recomputed to generate an empirical null distribution.

Adjusted group differences are reported with regression coefficients (β), robust t-statistics, permutation p-values, and covariate-adjusted Cohen’s d.

### Machine learning classification with LASSO

We trained a LASSO-regularized logistic regression classifier using EEG-derived biomarkers (BiS, E/I_HLP_, fE/I, E+I_HLS_, LZC) plus age and sex. The main model excluded DFA and γ-band features due to reliability concerns, achieving robust discrimination between MDD and HC (Fig. 5A). Additional analyses including DFA and γ-band predictors are presented in the Supplemental Information.

### Software and Reproducibility

All analyses were implemented in MATLAB (Statistics and Machine Learning Toolbox) and Python. Random seeds were fixed (rng(1)).

Details of preprocessing, metric definitions, and full per-band statistical results are provided in the Supplemental Information.

## Results

### Frequency-specific criticality in MDD

To test the hypothesis that HC brains operate closer to criticality than brains of patients with MDD, we performed robust regression analyses of frequency-specific markers of brain criticality, complemented by simple group comparisons. For BiS (Figure 2A), covariate-adjusted regressions controlling for age and sex revealed significantly lower values in MDD than HC across θ, α, β, and γ bands, indicating reduced bistability and weaker stability of oscillatory states (Fig. 2Ab). The corresponding results provide an intuitive display of simple group differences that parallel the regression results (Fig. 2Aa). For DFA, robust regressions again showed diminished long-range temporal correlations in MDD, with the strongest effect in the α band (Fig. 2Bb); Welch comparisons (Fig. 2Ba) illustrate the same direction of effect. Although DFA estimates warrant caution given the relatively short data segments, both metrics converge on the conclusion that MDD brains deviate further from the critical regime than HC. Full per-band statistics (group means ±SE, 95% confidence intervals, Cohen’s d effect sizes, p-values, and Benjamini– Hochberg FDR–adjusted p-values) are provided in Supplemental Tables 2–11. Taken together, these findings indicate a frequency-specific yet consistent divergence from criticality in MDD. This divergence indicates that MDD is marked by frequency-specific shifts in emergent dynamics which may serve as candidate neurophysiological biomarkers.

**Figure 2.**
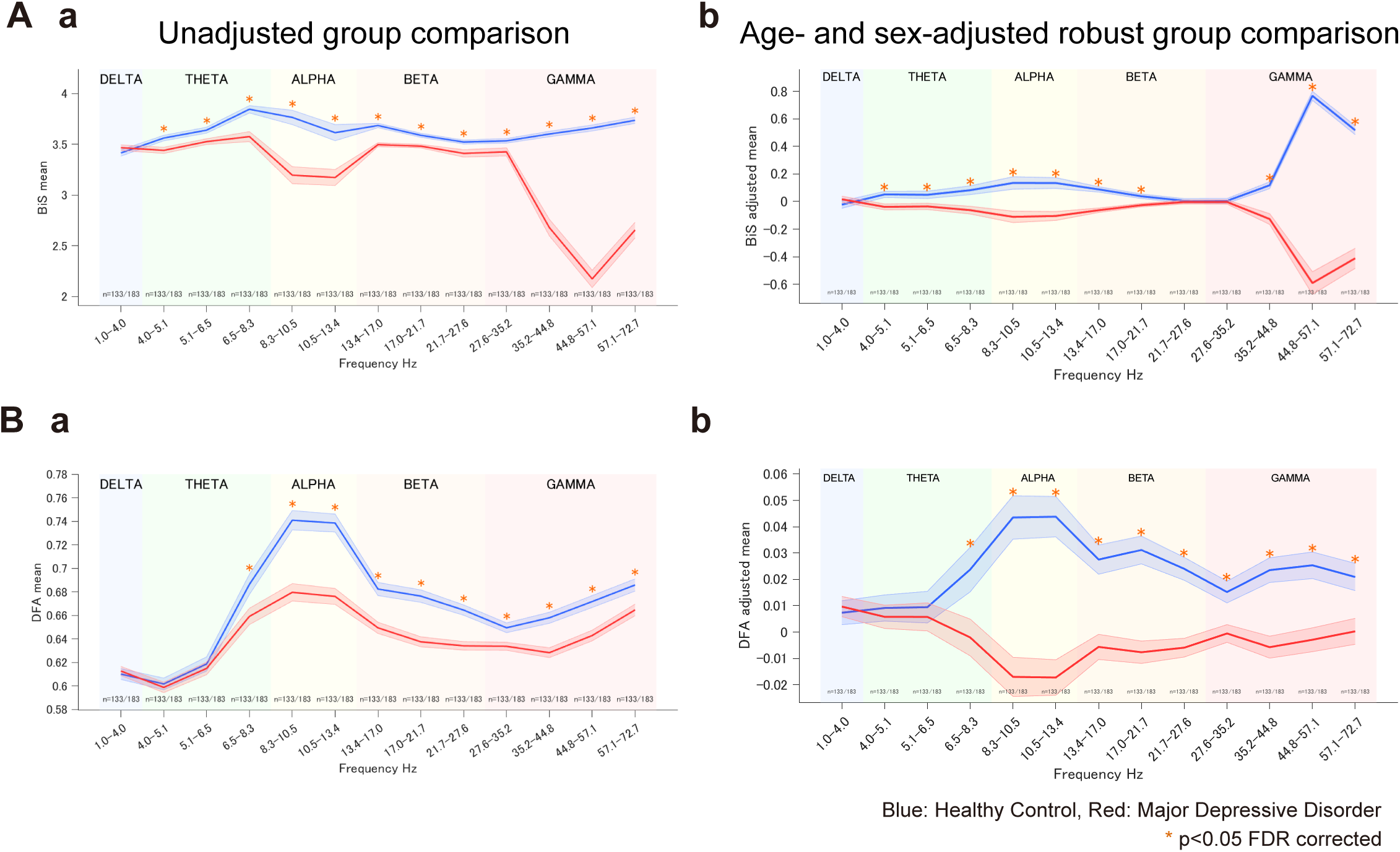
Group differences (HC vs MDD) in brain criticality: bistability (BiS) and detrended fluctuation analysis (DFA). (A) Bistability Index (BiS). (Aa) Unadjusted group means ± SE (channel-averaged per subject) with Welch tests. (Ab) Age- and sex-adjusted robust regression reveals lower BiS in MDD than in HC across θ, α, β, and γ, consistent with the raw contrasts. (B) Detrended Fluctuation Analysis (DFA). (Ba) Unadjusted group comparisons. (Bb) Age- and sex-adjusted robust regression reveals lower DFA in MDD compared to HC across è, á, â, and ã. Asterisks denote significance (black: p<.05 uncorrected; orange: FDR q<.05 across 13 bands). Numbers beneath the x-axis indicate per-band sample sizes (n=HC/MDD). Adjusted means in (Ab, Bb) are covariate-residualized estimates. Blue = HC; Red = MDD.

### Altered E–I coupling, with excess excitation in slower rhythms in MDD

We next examined indices of excitation–inhibition (E/I) balance and combined excitatory–inhibitory (E+I) strength. For the high-to-low power ratio (E/I_HLP_; Figure 3A), robust regressions revealed significantly higher values in MDD than HC in the θ and β bands, and significantly lower values in the γ band (Fig. 3Ab). Welch tests showed the same direction of effects in adjusted data (Fig. 3Aa). For the functional E/I ratio (fE/I; Fig. 3B), robust regressions indicated significantly reduced values in MDD in the θ band and increased values in the γ band (Fig. 3Bb), consistent with the BiS findings, with Welch tests again showing the same direction of effects (Fig. 3Ba). Finally, for the high– low separation (E+I_HLS_; Fig. 3C), which reflects the combined strength of excitatory and inhibitory inputs, robust regressions showed decreased values in MDD from θ through β bands, but increased values in δ and γ (Fig. 3Cb); group comparisons of unadjusted data again reveal a similar direction (Fig. 3Ca). Full per-band statistics (mean differences, effect sizes, and corrected p-values) are provided in Supplemental Tables 2–11. Collectively, these results indicate that MDD is characterized by frequency-specific alterations of both E/I balance and combined E+I strength. Furthermore, group effects on E/I_HLP_ and E+I_HLS_ together indicate a higher E/I ratio and lower combined E+I, implying decreased inhibitory drive underlying θ oscillations in MDD. Similarly, lower E/I and higher combined E+I indicate increased inhibitory drive underlying γ oscillations in MDD. Considering potential for high-frequency EEG activity susceptibility to muscle artifacts, we interpret γ-band effects with caution. Nonetheless, these frequency-dependent alterations of E/I dynamics represent a neurophysiological fingerprint of MDD.

**Figure 3.**
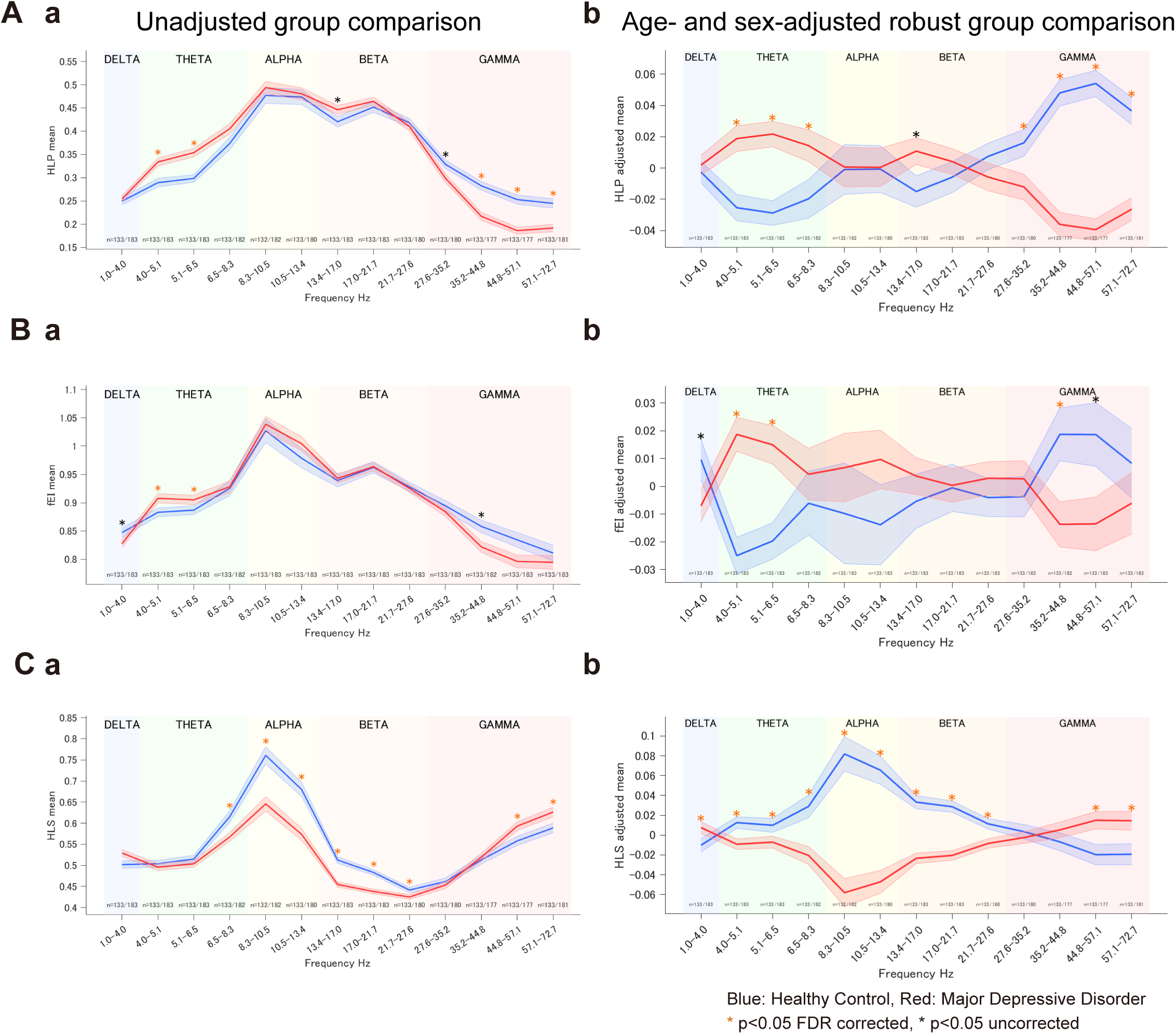
Group differences (HC vs MDD) in excitation–inhibition (E/I) balance and E+I strength. (A) High-to-low power ratio (E/I_HLP_) as E-I balance index. (Aa) Unadjusted group means ± SE (channel-averaged per subject) with Welch tests. (Ab) Age- and sex-adjusted robust regression shows MDD > HC in θ and β, and MDD < HC in γ. (B) Functional E/I ratio (fE/I). (Ba) Unadjusted comparisons. (Bb) Adjusted robust regression shows MDD > HC in θ and MDD < HC in γ. (C) High–low separation (E+I_HLS_) as E+I strength index. (Ca) Unadjusted comparisons. (Cb) Adjusted robust regression shows MDD < HC from θ through β, and MDD > HC in δ and γ. Asterisks denote significance (orange: FDR q<.05 across 13 bands; black: p<.05 uncorrected). Numbers beneath the x-axis indicate per-band sample sizes (n=HC/MDD). Adjusted means in (Ab, Bb, Cb) are covariate-residualized estimates from the robust model. Blue = HC; Red = MDD.

### Null results in signal irregularity comparison

To test the hypothesis that signal complexity is reduced in MDD, we analyzed Lempel– Ziv complexity (LZC) using covariate-adjusted robust regression and complementary comparisons of direct group differences. The robust model controlling for age and sex showed no significant group effect (Fig. 4Ab), corroborated by a residual-based permutation test (Fig. 4Ac) and by largely overlapping residual distributions (Fig. 4Ad). Unadjusted Welch and Mann–Whitney tests likewise indicated no HC–MDD difference (Fig. 4Aa). Effect sizes were small in magnitude (|d| ≈ 0.1). These convergent results provide no evidence for altered LZC in MDD, suggesting that the broadband LZC does not differentiate groups in our dataset.

**Figure 4.**
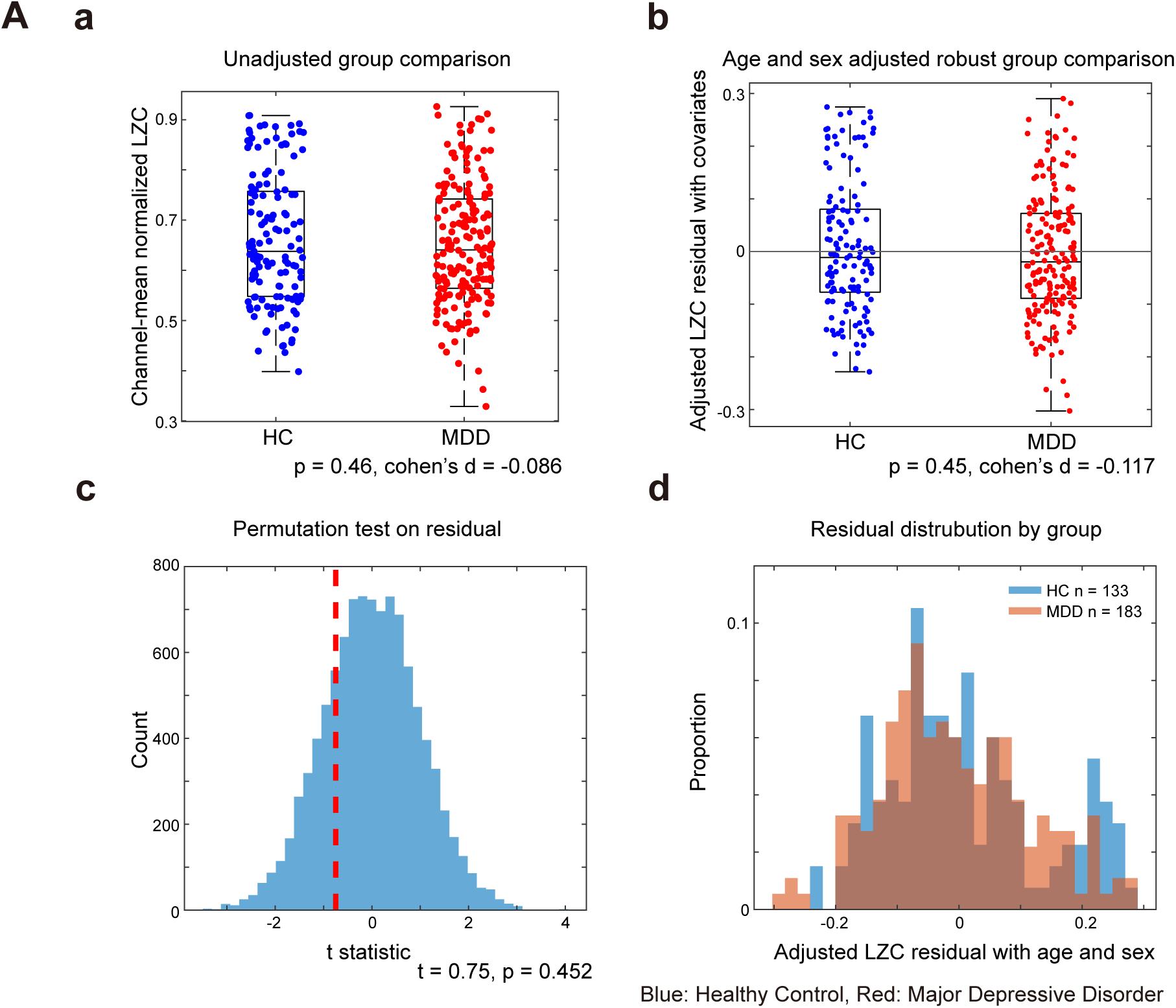
Group differences (HC vs MDD) in signal irregularity: Lempel–Ziv complexity (LZC) (a) Unadjusted group comparison (Welch t) shows no difference (p = .46, d = −0.09). (b) Covariate-adjusted robust comparison (age, sex) likewise shows no group effect (p = .45, d_adjusted_ = −0.12). (c) Freedman–Lane permutation of reduced-model residuals confirms the null (p_permutation_ = .45). (d) Age/sex-adjusted residual distributions overlap. LZC does not differentiate MDD from HC.

### Machine learning classification of HC vs MDD

We trained sparse logistic classifiers using the least absolute shrinkage and selection operator (LASSO) to test whether the multivariate EEG feature set separates patients with MDD from HCs. Models were trained on a stratified 70/30 split, with features z-scored by training statistics, and performance evaluated on the held-out test set using ROC and precision–recall (PR) curves (Fig. 5). Medium and higher γ-band dynamics were excluded due to potential contamination by muscle artifacts (27.6–35.2 Hz were retained). To be conservative, DFA features were omitted in the initial analysis over concerns about the length of available data segments. However, a comprehensive analysis, including DFA measures, was included in the Supplement (Supplemental Figure 3Aa).

**Figure 5.**
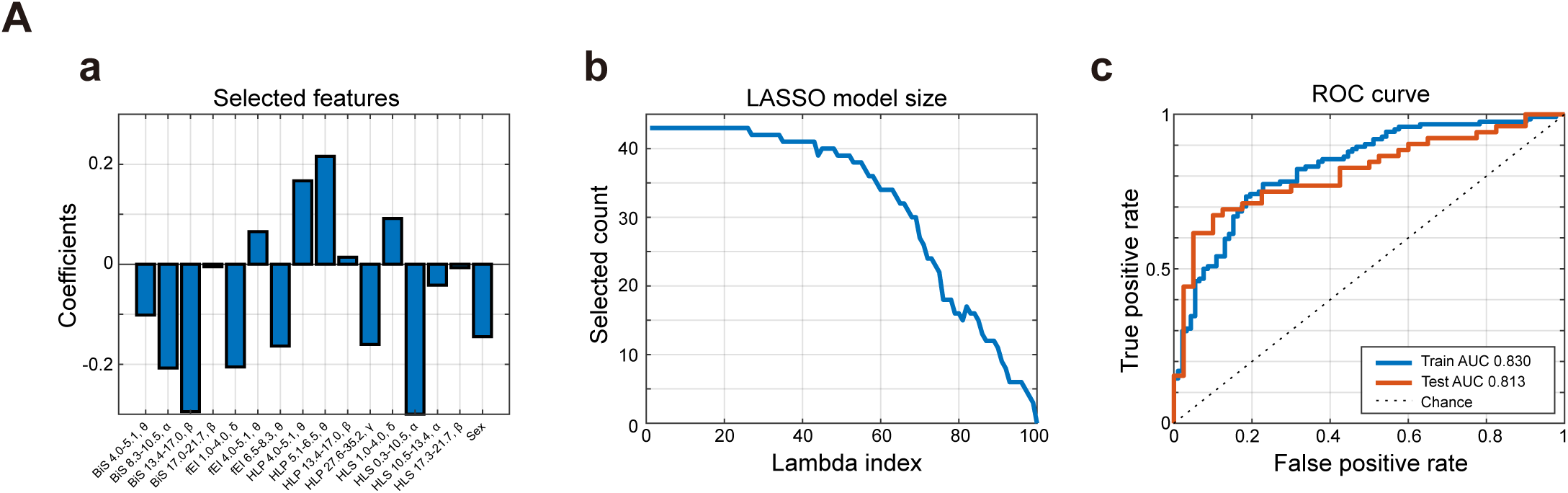
Machine learning classification of MDD versus HC. (A) LASSO logistic regression excluding DFA and high-ã features. (Aa) Non-zero coefficients show main contributions from E/I_HLP_, E+I_HLS_, BiS, and fE/I. (Ab) Path plot indicates sparsity across λ values. (Ac) ROC curves demonstrate high discrimination in both the training and test sets. The weight profile highlights frequency-specific E/I and criticality features as key predictive biomarkers of MDD.

### LASSO classification without DFA

At the optimal λ determined by 10-fold cross-validation (“1-SE rule”),, the classifier retained 16 features spanning markers of criticality (BiS), excitation–inhibition balance (fE/I and E/I_HLP_), excitation–inhibition strength (E+I_HLS_), and demographic covariates (age and sex). The 1-SE rule selects the simplest model whose cross-validation error is within one standard error of the minimum, favoring parsimony without loss of accuracy.

The largest positive weights were on E/I_HLP_ (5.1–6.5 Hz: +0.216, 4.0-5.1 Hz: +0.167), with additional smaller positives on E+I_HLS_ 1.0-4.0 Hz and fE/I 4.0-5.1 Hz; the largest negative weights were on E+I_HLS_ and BiS (8.3–10.5 Hz: −0.300, 13.4–17.0 Hz: −0.295, 8.3–10.5 Hz: −0.208), alongside fE/I (1.0-4.0 Hz: −0.205) and E/I_HLP_ (27.6–35.2 Hz: −0.160), with sex (female: −0.145). Overall, the weight profile is dominated by E/I_HLP_ and E+I_HLS_, with BiS providing additional (negative) contributions and fE/I showing mixed signs, consistent with frequency-specific E/I alterations and reduced bistability in MDD.

Several fE/I features contributed with both positive and negative weights, highlighting frequency-specific changes in E/I balance. On the training set, the model achieved AUC = 0.83 and accuracy = 0.75. On the independent test set, performance remained above chance with AUC = 0.81, accuracy = 0.74, sensitivity = 0.75, and specificity = 0.73. The path plot (Fig. 5Ab) demonstrated progressive sparsity with increasing λ, and coefficient visualization (Fig. 5Aa) revealed consistent contributions from criticality, E/I balance, and E+I strength indices. ROC curves (Fig. 5Ac) illustrated reliable and good discrimination against MDD versus HC.

These findings indicate that frequency-resolved measures of E/I balance and criticality (namely, bistability, but see also similar DFA effects in Supplementary Figure 1A) jointly contribute to separating MDD from HC. The weight profile was dominated by E/I_HLP_ and E+I_HLS_ features, with BiS providing additional contributions and fE/I showing mixed signs. This multivariate signature parallels the univariate results, showing altered network stability and E/I dynamics in depression, and demonstrating that the same physiological axes distinguishing groups at the feature level also carry predictive information at the multivariate level.

### LASSO classification with DFA

Including DFA features did not improve classifier performance. The DFA-inclusive model showed weaker generalization (test AUC = 0.75) compared to the DFA-excluded model.

### LASSO classification (full γ-inclusive)

When full γ-band features were included, the LASSO model achieved high accuracy (test AUC = 0.95) with predictors dominated by γ band BiS and E+I_HLS_. Because high-frequency EEG is prone to muscle artifacts, these γ-inclusive results are reported only in Supplemental Information.

## Discussion

### Summary

We hypothesized that major depressive disorder (MDD) brains diverge from criticality, because aberrant excitation-inhibition balance. The results supported these hypotheses. MDD brains diverged from criticality, with the strongest effects in the θ, α, β, and γ bands, using the bistability index (BiS) and detrended fluctuation analysis (DFA). We also found that, relative to HC, MDD brains showed decreased inhibitory drive in the θ band and increased inhibition in the γ band, as indicated by the high-to-low power ratio (E/I_HLP_), high-to-low power separation (E/I_HLS_), and the functional E/I ratio (fE/I). Going beyond prior work, our specific inferences about excitatory and inhibitory strength (as opposed to, more simply E/I balance) were made possible by the recent high-to-low-power oscillatory index (E+I_HLS_), which is quantified by the separation of high- and low-power oscillations (_HLS_). Here we found that (E/I_HLS_) was decreased from the θ through β bands but increased in the γ band. These results suggest that MDD is not uniformly shifted toward excitation or inhibition but instead shows frequency-specific deviations in both directions, as well as the alteration in the excitation–inhibition strength. In contrast, irregularity measured by Lempel–Ziv complexity (LZC) showed no significant differences between groups, whether or not we adjusted for age and sex, indicating that broadband irregularity may not be a biomarker of MDD.

Finally, even conservatively excluding γ-band and DFA features, the machine learning method with LASSO regression classified MDD from HC with features of criticality, E/I balance, and E+I strength dynamics (BiS, E/I_HLP_, E+I_HLS_, and fE/I). Collectively, these findings demonstrate that criticality, E/I balance, and E+I strength are significant physiological signatures of MDD as a complex system fingerprint.

### Interpretation

Our findings indicate that MDD is characterized by a departure from the critical operating regime of large-scale neural dynamics, expressed as frequency-specific disturbances of the E/I balance, decreased inhibition in θ and excess inhibition in γ. Also, a shift away from criticality implies suboptimal information transmission and reduced dynamic range, which we speculate may constitute a neural substrate for both mood dysregulation and cognitive impairment in MDD.

Band–specific E/I patterns are also a novel finding, variously consistent with prior postmortem and MRS evidence of regionally differentiated neurotransmission (reduced GABAergic tone and altered glutamatergic signaling) (Yuksel and Oenguer 2010, Godfrey, Gardner et al. 2018, Hu, Tan et al. 2023). For example, the default mode network (DMN) tends toward relative over-excitation, while the frontoparietal control regions tend toward relative over-inhibition (Hu, Tan et al. 2023). Regarding broadband decreases in critical dynamics (bistability and long-range temporal correlations), we speculate that a Griffiths phase, which normally expands the critical regime over a range of E/I values (Moretti and Munoz 2013), shrinks in MDD due to mechanism-specific alterations in E/I balance across brain systems, restricting emergent dynamics, on average. Shrinkage of Griffiths phases undermines the extended regime that normally supports adaptive responses, suggesting that self-organizing mechanisms in neural systems may fail to operate as robustly in MDD.

The neural basis of θ oscillations involves both hippocampal and cortical generators, paced by projections from septal cholinergic and GABAergic neurons (Vertes and Kocsis 1997, Buzsáki 2002). In human EEG, the fronto-midline θ rhythm has been linked to cognitive control processes (Cavanagh and Frank 2014) and may contribute to the cognitive deficits observed in patients with MDD (Pizzagalli 2011). The neural basis for γ oscillations primarily relies on parvalbumin-positive GABAergic fast-spiking interneurons, and is present in both the neocortex and hippocampus (Cardin, Carlén et al. 2009, Sohal, Zhang et al. 2009). They are distributed throughout the neocortex, including the frontoparietal regions. We speculate that an impairment in γ oscillations in frontal and parietal cortex may be related to depression symptoms, and cognitive deficits in MDD. Further, θ–γ coupling constitutes a key neural mechanism supporting diverse cognitive functions, including consciousness, attention, and working memory (Ursino and Pirazzini 2024). By coordinating slow θ and fast γ oscillations, this coupling provides a temporal coding scheme that enables information processing and integration across distributed brain networks (Canolty and Knight 2010, Lisman and Jensen 2013). Alterations in corresponding excitation-inhibition mechanisms may therefore play a central role in the pathophysiology of MDD.

Treatment studies also converge on the hypothesis of aberrant E/I balance in MDD. In responders to electroconvulsive therapy (ECT), EEG-based functional E/I ratios increased selectively in the β band (12–28 Hz), indicating a shift toward excitation (Stuiver, Pottkämper et al. 2023). Considering transcranial magnetic stimulation (TMS), whole-brain artificial neural network simulations of TMS–EEG effects suggested a shift toward inhibition in responders, accompanied by suppression of α and θ oscillations (Momi, Wang et al. 2025). Finally, accumulating evidence suggests that fast-acting anti-depressant medication have a common effect of increasing excitatory spine growth (Liao, Dua et al., 2025) – an effect which could plausibly offset the over-inhibition we infer with respect to γ-band oscillations. Across modalities, recovery appears to involve restoring the system toward criticality via calibrated E–I rebalancing.

The LASSO classification results highlight that MDD can be distinguished from HC not by a single dominant feature, but by a distributed set of biomarkers spanning criticality, E/I balance, and E+I strength. The sparsity of the retained predictors underscores that only a subset of frequency-specific measures carries the strongest discriminative power, with E/I_HLP_ and E+I_HLS_ dominating the weight profile. Importantly, BiS, E/I_HLP_, fE/I, and E+I_HLS_ contributed with positive and negative weights, suggesting that deviations from criticality and shifts in E/I balance and E+I strength manifest heterogeneously across frequency bands. This distributed pattern reinforces the notion that depression is not characterized by a uniform shift toward excitation or inhibition but instead reflects a complex reorganization of network stability and E/I dynamics. The alignment of the multivariate weights with the univariate group-level results strengthens the validity of these physiological axes as mechanistically meaningful markers, while also demonstrating their utility for predictive modeling. Together, these findings indicate that criticality-informed EEG metrics offer both explanatory power and diagnostic utility, supporting their potential as biomarkers for MDD.

### Considerations and limitations

Several limitations should be noted. First, DFA estimates were based on short data segments of 49 or 60 seconds and, therefore, excluded from machine learning analyses (reported in Supplemental Information). Second, γ-band metrics are prone to muscle artifacts and were treated as exploratory. Third, EEG data were acquired with different systems in HC and MDD groups (ANT vs. ActiCAP); harmonized preprocessing was applied, but residual effects cannot be fully ruled out. Fourth, group demographics were not perfectly matched on age and sex, although we addressed this using robust regression adjusted for these covariates. At the time of EEG acquisition, nearly all the MDD patients were concurrently taking various types of psychotropic medications which may impact the results. Future studies should evaluate medication effects apart from diagnostic category. Finally, our cross-sectional design precludes causal inference.

### Conclusion

Our large-scale EEG study demonstrates that MDD is characterized by deviations from brain criticality, with frequency-specific excitation–inhibition imbalance, and decreased inhibition in mechanisms operating in the θ-band and increased γ-band inhibition. Together, these alterations define a complex system fingerprint of MDD, highlighting their potential as a biomarker for psychiatric disorders more generally (O’Byrne and Jerbi, 2022; Hengen and Shew, 2025). By integrating metrics of criticality, E/I balance, and E+I strength, resting EEG may support both diagnostic classification and treatment monitoring, and ultimately guide personalized interventions aimed at restoring the brain toward criticality.

## Author Contributions

1. Conceptualization AM and AW
2. Data curation AM, FF, LC, and AW
3. Formal analysis AM and AW
4. Funding acquisition AW
5. Investigation AM, LC, and AW
6. Methodology AM, LP, and AW
7. Project administration AW
8. Resources AM and AW
9. Software AM and AW
10. Supervision AW
11. Validation AM, AB, and AW
12. Visualization AM
13. Writing – original draft AM
14. Writing – review & editing AM, LC, LP, and AW

## Lead contact

Further information and requests for resources should be directed to and will be fulfilled by the lead contact, Andrew Westbrook.

## Acknowledgements

This work was supported by Grant No. R00 MH125021 of NIMH to AW. We are grateful to Li Xin Lim (Rutgers), Arthur-Ervin Avramiea (CNCR), and Klaus Linkenkaer-Hansen (CNCR).

## Supplemental Information

### EEG acquisition (MDD)

Clinical EEG was collected at the Butler Hospital TMS Clinic using a 64-channel ANT Neuro system (EEGO software, 10–10 montage; CPz reference, AFz ground). Signals were recorded with eyes closed, and no online filters were applied. For the present study, we analyzed the pre-stimulation baseline obtained prior to the first TMS session of each patient’s earliest treatment series. The raw data were acquired at 2000–2048 Hz for 5–10 minutes and were subsequently resampled to 512 Hz and trimmed to ∼5 minutes of artifact-free resting EEG per subject by the data provider. For analysis, each ∼5-minute recording was then segmented into five contiguous 1-minute epochs and re-ordered in a fixed sequence (1–3–5–2–4) to construct the final time series, aligning the epoch structure with the 1-minute block design used for HC data. To maximize the number of patients included in the analysis, recordings shorter than 5 minutes were standardized to a common length of 4.65 minutes by removing 10.6 seconds from both the beginning and the end of each recording.

### EEG acquisition (HC)

Resting-state EEG was recorded in the LEMON study using a 62-channel active electrode cap (ActiCAP, Brain Products GmbH, Germany; 10–20 montage) in combination with two additional ECG channels (Babayan et al., 2019). Recordings alternated between eyes-open and eyes-closed conditions in 1-minute blocks for a total of 16 min, beginning and ending with eyes open. During eyes-open periods, participants fixated on a low-contrast cross presented on a gray background.

For the present study, we extracted only the eyes-closed segments to match the acquisition protocol used in the MDD patient sample. The 1-minute eyes-closed epochs were concatenated, down-sampled from the original 2,500 Hz to 512 Hz (to align with the patient recordings) and then trimmed to ∼5 minutes per subject. To match the data length used for the HC and MDD recordings, an additional 10.6 seconds were removed from both the beginning and the end of each HC recording, resulting in a length of 4.65 minutes after the preprocessing.

### Metrics

#### Bistability (BiS)

We estimated the probability density function (PDF) of narrow-band EEG power to quantify the bistability of neuronal oscillations. Bistability reflects the tendency of amplitude fluctuations to alternate between high- and low-power states, resulting in a bimodal distribution in critical systems (Freyer, Aquino et al. 2009).

For each channel × frequency band, the analytic amplitude A(t) was obtained using the Hilbert transform after bandpass filtering, squared to yield instantaneous power, and normalized by the median. The empirical PDF of A(t) was then estimated from histograms with 200 equally spaced bins.

To assess bimodality, we compared a unimodal exponential distribution against a bi-exponential mixture, both fitted by maximum likelihood estimation. Model evidence was evaluated using the Bayesian Information Criterion (BIC), and the Bistability Index (BiS) was defined as:

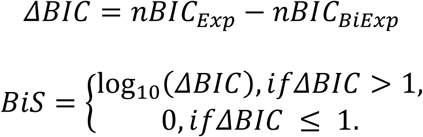

For the model fitting, parameters were optimized using the Nelder–Mead algorithm. The two exponents for the bi-exponential model were both initialized to the value estimated from the single-exponential fit, and the mixing weight was initialized to 0.5. During optimization, we enforced the constraints that the first exponent must be larger than the second and that the mixing weight must lie between 0 and 1.

Following prior work (Avramiea, Diachenko et al. 2025), our computation of nBiC treats every dataset (all of which contain n ≥140,000 samples), as though they contain 100,000 samples:

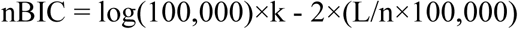

where k is the number of free parameters (k = 1 for the single-exponential model and k = 3 for the bi-exponential model) and L is the likelihood function. This normalization was implemented to maintain methodological consistency with prior work, although all EEG segments were of equal length (n ≈ 140,000 samples) and thus the correction has a negligible effect on the results.

#### Detrended Fluctuation Analysis (DFA)

We estimated Hurst exponents from band-limited amplitude envelopes of the EEG. Signals were band-pass filtered into 13 logarithmically spaced frequency bins between 1 and 80 Hz using a finite impulse response (FIR) filter designed with the window method (implemented in MNE). Filtering used a Hamming-window FIR filter, applied in zero-phase (bidirectional) mode, with transition bands determined automatically by MNE’s default heuristics. Window sizes were initially generated as 81 logarithmically spaced points between 0.1 and 1000 seconds (approximately 20 per decade) and converted to samples according to the sampling frequency. After rounding to integer samples and restricting the range to the fitting interval [*S_min_*, 30 s], approximately 50 distinct window sizes remained for DFA estimation. Windows were slid with 50% overlap. For each frequency bin, the DFA exponent was estimated by fitting a straight line to the log–log plot of fluctuation versus window size within the fitting interval. The upper bound of 30 seconds is conventional in long-range temporal correlation analyses of human resting EEG (Hardstone, Poil et al. 2012, Avramiea, Diachenko et al. 2025). The lower bound *S_min_* was band-specific and taken from a predefined schedule [5.0, 5.0, 5.0, 3.981, 3.162, 2.238, 1.412, 1.122, 0.794, 0.562, 0.398, 0.281, 0.141] seconds, matched to the lower edge of each frequency bin.

The DFA computation followed three standard steps:

##### 1. Signal profile construction

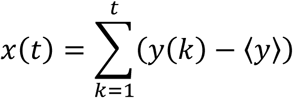

where y(k) is the demeaned signal, ⟨y⟩ denotes the temporal mean of the signal, and x(t) is the integrated profile.

##### 2. Local detrending and fluctuation

For each window of length *s*, a first-order polynomial *y*_*s*_(*k*) was fit by ordinary least squares to the profile *Y*(*k*) and subtracted. The fluctuation at scale *s* was then defined as the across-windows average of the within-window standard deviation:

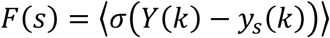

##### 3. Scaling law

Fluctuations across time scales were assessed by regressing *l*og*F*(*s*) against in the fitting interval, yielding the DFA (Hurst) exponent *α* from the slope:

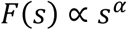

#### The proportion of high- and low-power oscillations (E/I_HLP_)

The bi-exponential model of the power distribution decomposes the signal into a low-power and a high-power component, which can be interpreted as reflecting inhibitory- and excitatory-dominated network states, respectively. Inhibition stabilizes the system and favors low-amplitude activity, whereas excitation promotes large-amplitude oscillations. Mixing weight δ represents the relative contribution of the two exponential components describing low- and high-power oscillatory states. The model is constrained such that the first exponent corresponds to the low-power state and the second to the high-power state. Thus, δ can be interpreted as the proportion of high-power activity. δ values near 0 indicate predominantly low-power (inhibitory) states, values near 1 indicate predominantly high-power (excitatory) states, and δ values around 0.5 indicate a balanced contribution of both. δ was only interpreted when the signal showed clear bistability (BiS ≥ 2.5) and δ fell within the range 0.02–0.98; otherwise, it was set to NaN.

To ensure interpretability, E/I_HLP_ was only computed when the signal exhibited sufficient bistability (BiS ≥ 2.5) and when δ lay within the range 0.02–0.98; otherwise, E/I_HLP_ values were set to NaN. Under these conditions, δ provides an index of the proportion of high- to low-power oscillations and serves as a proxy for the excitation– inhibition balance of the underlying network. We applied a threshold of BiS ≥ 2.5, under which the δ–BiS relationship becomes stable and δ values closely match independent estimates of excitation–inhibition balance (Avramiea, Diachenko et al. 2025).

#### Functional E/I ratio (fE/I)

The functional excitation–inhibition ratio (fE/I) provides a model-based estimate of cortical excitation–inhibition balance by quantifying the coupling between oscillatory amplitude and long-range temporal correlations (LRTC) (Bruining, Hardstone et al. 2020). Excitatory activity tends to increase oscillatory amplitude, whereas LRTC increase with excitation up to the critical point and become negatively correlated with excitation beyond that. Thus, the correlation between amplitude and fluctuation scaling provides a proxy for the underlying excitation–inhibition ratio.

##### 1. Band-pass filtering and amplitude extraction

EEG signals were first band-pass filtered into the predefined frequency bins using a finite impulse response (FIR) filter (Hamming window, zero-phase, bidirectional). The analytic amplitude Y(t) was then extracted as the absolute value of the Hilbert transform of the filtered signal.

##### 2. Signal profile construction

For each channel × frequency bin, the amplitude envelope was demeaned and cumulatively summed to generate a signal profile:

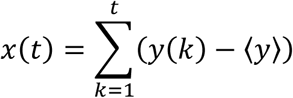

where y(k) is the instantaneous amplitude, ⟨y⟩ its temporal mean, and x(t) the integrated profile.

##### 3. Windowing and amplitude normalization

The signal profile was segmented into 5-second windows with 80% overlap. Within each window, the profile was divided by the mean amplitude of that window. This normalization step removes scale differences in absolute oscillatory power.

##### 4. Local detrending

Within each normalized window, a first-order polynomial *y*_*s*_(*k*) was fit by least squares and subtracted from the profile, yielding detrended, amplitude-normalized windows.

##### 5. Fluctuation function

For each window, the standard deviation of the detrended, normalized profile was computed to yield the normalized fluctuation function:

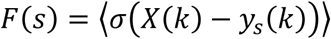

##### 6. Correlation with amplitude

Across windows, we computed the Pearson correlation coefficient r between the windowed mean amplitude and the windowed normalized fluctuation F(s). The fE/I ratio was then defined as:

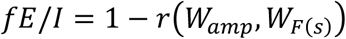

Where *W*_*amp*_ denotes windowed mean amplitudes and *W*_*F*(*s*)_ the corresponding windowed fluctuations. By construction, *fE* / *I* ≈ 1 indicates balanced excitation– inhibition dynamics, whereas values above or below 1 reflect shifts toward excitation- or inhibition-dominated regimes, respectively.

##### 7. Quality control

Following prior work, the fE/I estimate was set to NaN when the DFA exponent of the underlying signal profile did not exceed a minimum threshold of 0.6, ensuring that fE/I was only computed in signals with sufficient long-range temporal correlations.

### Excitation-inhibition strength

#### Separation of High- and Low- Power Oscillations (E+I_HLS_)

We quantified how separated high- and low-amplitude states were in the power distribution – an index of combined excitatory and inhibitory strength (E+I; Avramiea, Diachenko et al., 2025). For each bi-exponential fit, we identified the location of the low-power and high-power peaks and computed their distance on a logarithmic scale:

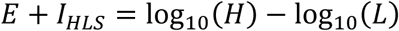

Where L and H denote the positions of the low- and high-power peaks, respectively. This measure reflects the order-of-magnitude gap between the two states, with larger values indicating a clearer distinction between low- and high-amplitude oscillatory regimes.

To ensure reliable interpretation, E+I_HLS_ was only calculated when the signal showed sufficient bistability (BiS ≥ 2.5) and when both states contributed substantially to the distribution (0.02 ≤ δ ≤ 0.98). In all other cases, E+I_HLS_ values were set to NaN.

#### Lempel–Ziv complexity (LZC)

We estimated signal irregularity by computing Lempel–Ziv complexity (LZC) using a single-pass Lempel–Ziv–Welch (LZW) dictionary-building procedure in MATLAB 2020a (Comsa 2019). Starting from an initial dictionary containing the two symbols [0, 1] the binary sequence was scanned left-to-right. At each step, the current phrase was extended by one symbol; when the extended phrase was not found in the dictionary, it was added as a new entry, and a new phrase was started. The complexity of the sequence was defined as the final dictionary size, representing the number of unique patterns identified. To account for sequence length and symbol frequency, we normalized this value by dividing it by the mean complexity obtained from ten randomly shuffled versions of the same sequence (N=10). Larger normalized values indicate more irregular or less structured signals. We note that the implementation uses random shuffling without an explicitly fixed seed; thus, normalized values may vary minimally across runs.

Although the ∼4.65-min time series was assembled from five separate segments (49–60–60–60–49 s) rather than a single continuous recording, this does not compromise the reliability of the LZC estimates. Rivolta et al. demonstrated that as few as 1,000 datapoints are sufficient for stable LZC estimation during sleep (Rivolta, Migliorini et al. 2014), and each of our segments greatly exceeded this length.

### Statistical analysis for BiS, DFA, fE/I, E/I_HLP_, and E+I_HLS_

For E/I_HLP_ and E+I_HLS_, channel-level estimates returned some NaN’s as described above. At the subject level, we averaged across channels, omitting NaNs, such that values were computed from available channels only. If all channels were missing for a given subject and band, the subject was excluded from subsequent group-level analyses.

### Machine learning classification with LASSO

We trained a LASSO-regularized logistic regression classifier to distinguish MDD from HC. The feature matrix comprised EEG-derived biomarkers indexing criticality (BiS), excitation–inhibition balance (E/I_HLP_, fE/I), excitation-inhibition strength (E+I_HLS_), and complexity (LZC), with age and sex added as covariates. To minimize contamination by potential high-γ artifacts, we excluded bands 11, 12, and 13 (35.2-44.8 Hz, 44.8-57.1 Hz, and 57.1-72.7 Hz, respectively) for BiS, fE/I, E/I_HLP_, and E+I_HLS_. Rows containing missing (NaN) or non-finite (Inf) values were excluded. Participants (183 MDD, 133 HC) were split into stratified training (70%) and test (30%) sets with a fixed random seed, ensuring balanced class ratios. Features were standardized to zero mean and unit variance using training-set statistics, with the same parameters applied to test data to prevent data leakage.

For the classifier, 100 candidate regularization parameters (λ) were evaluated by 10-fold cross-validation within the training set. The optimal λ was selected according to the one-standard-error rule, yielding a sparse and interpretable model. Non-zero coefficients at this λ were retained as predictive features, and an unregularized logistic regression model was refit using only these features on the training data and then applied to the held-out test set. Predicted probabilities of MDD were converted to class labels (MDD if ≥0.5, HC if <0.5). Model performance was quantified by area under the receiver operating characteristic curve (AUC), accuracy, sensitivity, and specificity.

Figure 5A illustrates (a) non-zero model coefficients, (b) the number of features selected across λ values (regularization path), and (c) ROC curves for training and test sets.

We also evaluated models including DFA and γ-band features. Because our EEG data lengths were shorter than the traditional length recommended for reliable DFA estimation, and γ-band activity may partly reflect muscle noise, these measures were not included in the main analysis (Figure 5A). For completeness, Supplemental Figure 3A shows classification performance when DFA was added, and Supplemental Information reports results from models including both DFA and γ-band features.

## Supplemental Results

### LASSO classification with DFA

We next examined whether adding DFA features altered classifier performance. At the optimal λ (10-fold CV, 1SE rule), the model retained features, including markers of criticality (BiS, DFA), E–I balance (fE/I, E/I_HLP_), E–I strength (E+I_HLS_), age, and sex. The coefficient map showed the largest positive weights on E/I_HLP_ (hlp b 02 (θ: 4.0-5.1 Hz) = +0.194, hlp b 03 (θ: 3.1-6.5 Hz) = +0.121) and fE/I (fei b 02 (θ: 4.0-5.1 Hz) = +0.133), whereas the largest negative weights were on fE/I (fei b 01 (δ: 1.0-4.0 Hz) = −0.382), E+I_HLS_ (hls b 05 (α: 8.3-10.5 Hz) = −0.193, hls b 08 (β: 17.0-21.7 Hz) = −0.121), BiS (bis b 07 (β: 13.4-17.0 Hz) = −0.182, bis b 05 (α: 8.3-10.5 Hz) = −0.065), and DFA (dfa b 05 (α : 8.3-10.5 Hz) = −0.128, dfa b 06 (α: 10.5-13.4 Hz) = −0.009) (Supp. Fig. 5Aa). The path plot illustrates progressive sparsity as λ increases, with the number of non-zero coefficients steadily decreasing (Supp. Fig. 3Ab). At the optimal λ determined by cross-validation, the model retained 13 features. ROC curves indicated above-chance, but weaker discrimination compared with the DFA-excluded model: training AUC = 0.84 (ACC = 0.76), test AUC = 0.75 (ACC = 0.68, sensitivity = 0.77, specificity = 0.56) (Supp. Fig. 3Ac).

Together, these results show that while DFA features were selected by the model, their inclusion did not improve generalization, and test performance remained lower than the model without DFA.

### LASSO classification with full γ-bands

Including γ-band features (bands 11–13) yielded a LASSO model that retained 12 predictors, with the coefficient map (Supp. Fig. 3Ba) dominated by large γ-band BiS and E+I_HLS_ weights (bis b 12 (γ: 44.8-57.1 Hz) = −2.42, hls b 13 (γ: 57.1-72.7 Hz) = +0.80). The path plot showed a sharp drop in the number of selected coefficients near λ ≈ 0, followed by a more gradual decline as λ increased (Supp. Fig. 3Bb). At the cross-validated optimal λ, 12 predictors were retained. The ROC curve (Supp. Fig. 3Bc) indicated excellent performance (train AUC = 0.99, test AUC = 0.95; ACC = 0.90, sensitivity = 0.90, specificity = 0.90). However, the coefficient map (Supp. Fig. 3Ba) was dominated by γ-band features, with BiS in band 12 showing the largest weight (β = −2.42). These features likely contributed substantially to the improved discrimination. However, because high-frequency EEG is particularly susceptible to muscle artifacts, the γ-inclusive findings should be interpreted with caution and are reported only as supplemental.

**Supplemental Figure 1.**
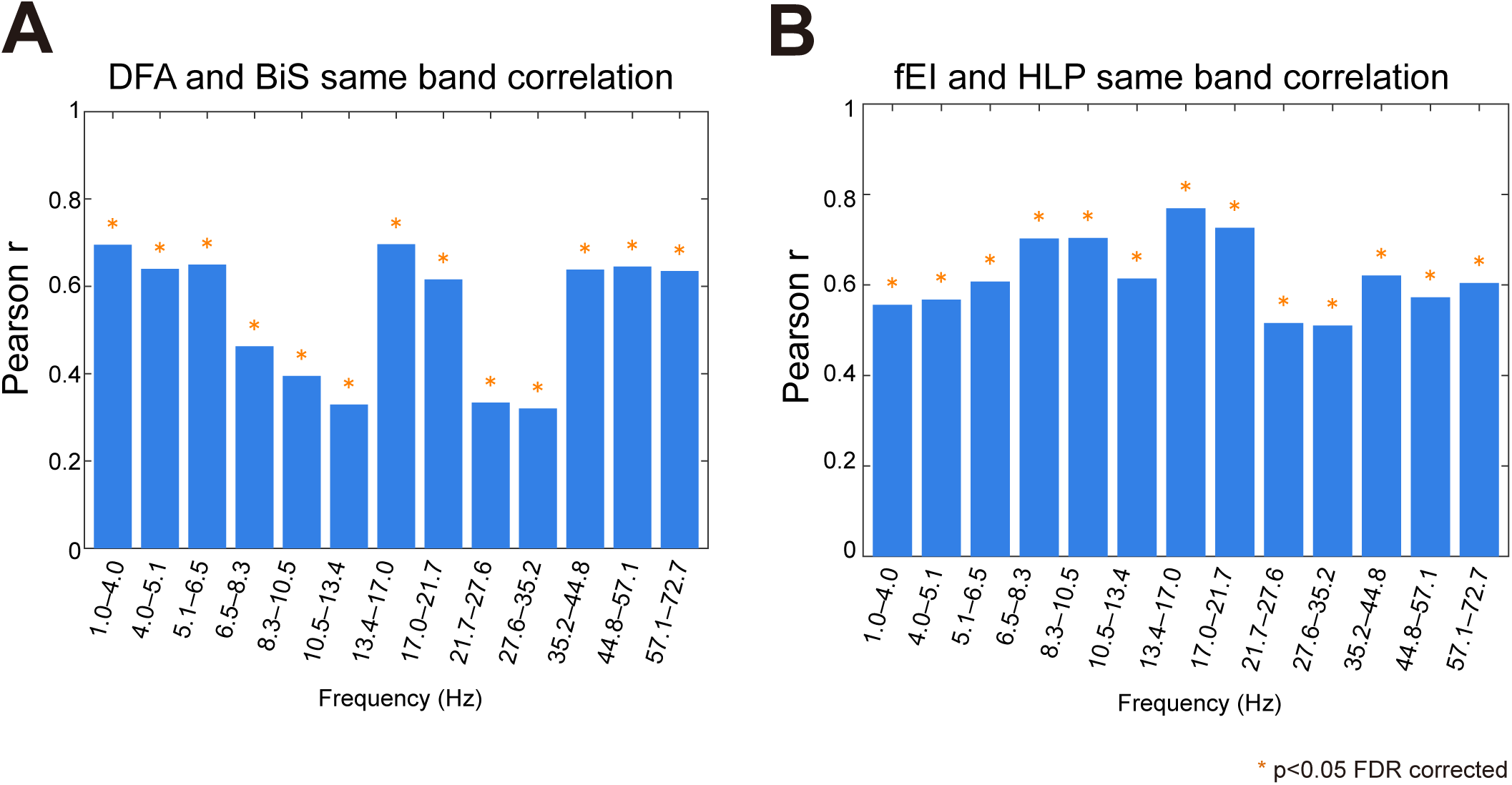
Same-band correlations between metrics. (A) Pearson correlation coefficients (r) between detrended fluctuation analysis (DFA) and bistability (BiS) across frequency bands. (B) Pearson correlation coefficients (r) between functional excitation–inhibition ratio (fEI) and high-to-low power ratio (HLP) across frequency bands. Orange asterisks indicate statistical significance (* p < 0.05, FDR corrected for 13 frequency bands). All available participants (n = 316) were included in the analyses, with missing values treated as NaNs and skipped in pairwise correlations.

**Supplemental Figure 2.**
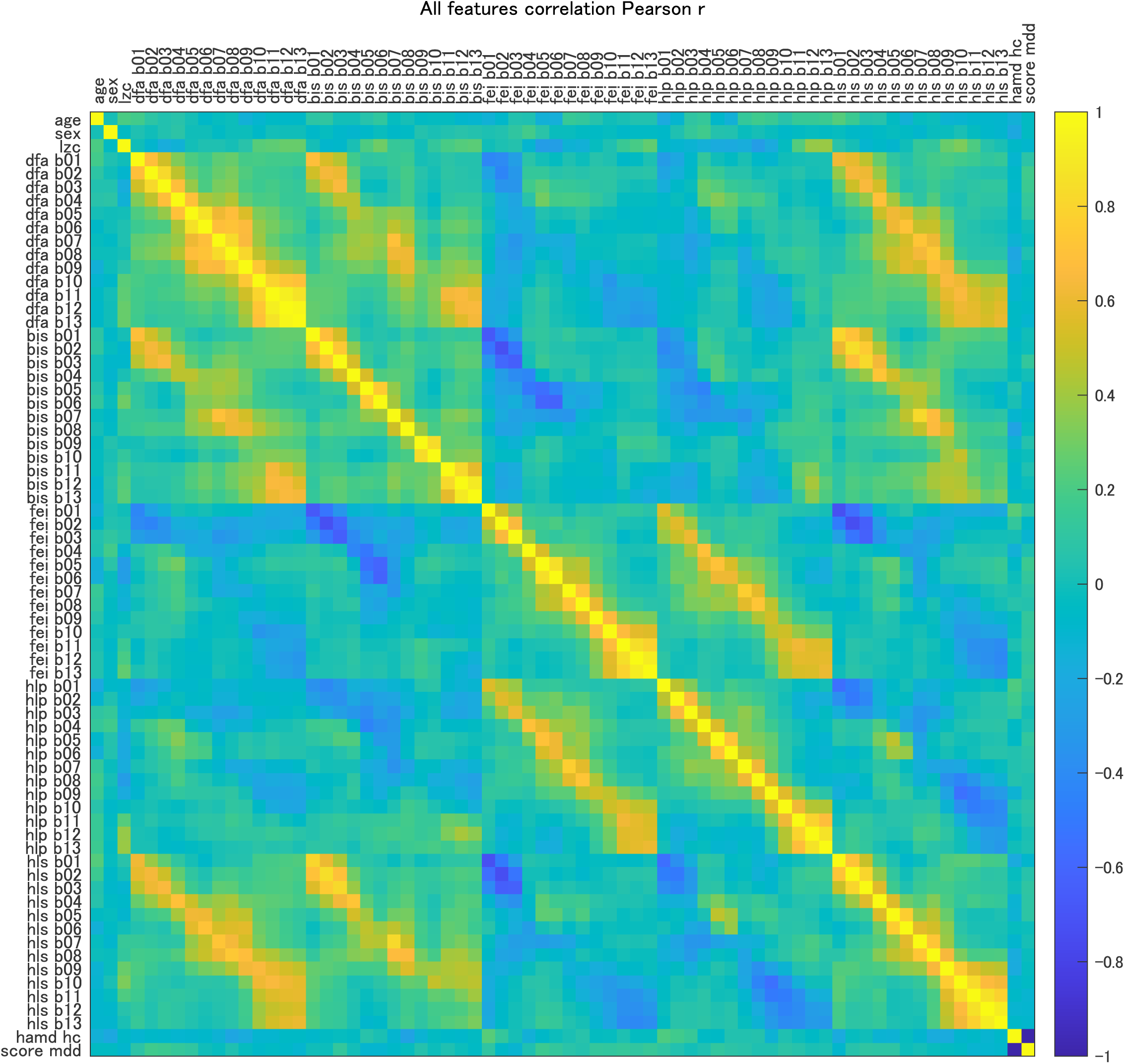
Feature-to-feature correlation matrix. Values represent the Pearson correlation coefficient between pairs of extracted features, age, sex, and depression scores (HAMD for HC ≤(mamd hc), IDS-SR for MDD (mdd score)). The right color scale indicates the correlation strength (-1 ≤ r ≤ 1). Warm colors (yellow) indicate positive correlations, and cool colors (blue) indicate negative correlations. Frequency bands b01–b13 correspond to 13 logarithmically spaced sub-bands.

**Supplemental Figure 3.**
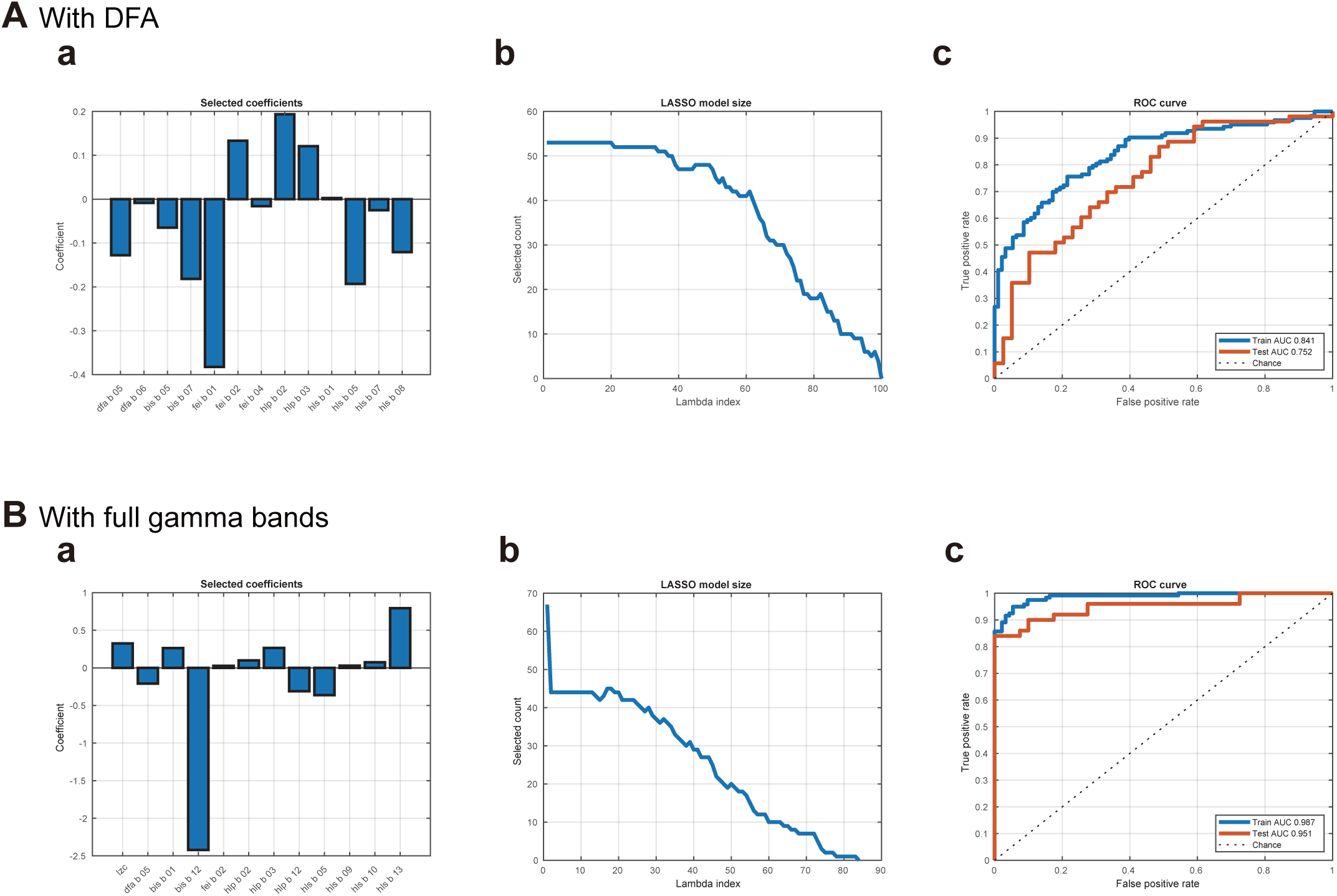
Machine learning classification with and without DFA and full gamma bands. (A) With DFA included. (Aa) LASSO logistic regression coefficients for the optimal model. Frequency bands b01–b13 correspond to 13 logarithmically spaced sub-bands. (Ab) Path plot indicates model sparsity across λ values (model size as a function of the regularization parameter). (Ac) Receiver operating characteristic (ROC) curves demonstrate classification performance in the training and test sets (Train AUC = 0.841, Test AUC = 0.752). (B) With full gamma bands included. (Ba) LASSO coefficients for the optimal model reveal additional contributions from high-γ features. (Bb) Path plot indicates sparsity across λ values. (Bc) ROC curves show improved discrimination (Train AUC = 0.987, Test AUC = 0.951).

**Supplemental Table 1.**
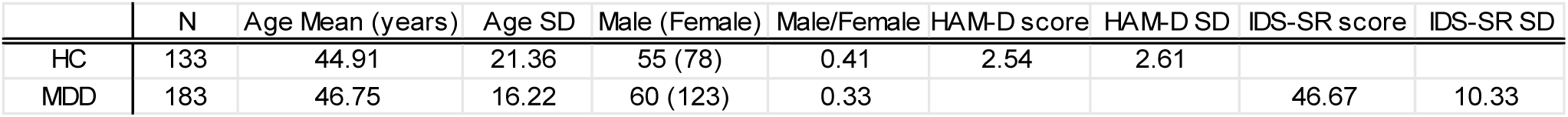
Demographics and clinical scores for healthy controls (HC) and patients with major depressive disorder (MDD). Values are mean ± SD unless noted. Sample sizes: HC N=133; MDD N=183. Age: HC 44.91 ± 21.36 years (HC age was recorded in 5-year bins; mean and SD were estimated from bin midpoints); MDD 46.75 ± 16.22 years; between-group difference not significant (Welch t = −0.84, p = 0.404; 95% CI −6.19 to 2.50). Sex: HC 55 male and 78 female (male/female ratio 0.41); MDD 60 male and 123 female (male/female ratio 0.33); distribution not significantly different between groups (chi-square = 2.44, df = 1, p = 0.118; Cramer’s V = 0.088). Depression measures are group-specific and cannot be compared across groups: HC, Hamilton Depression Rating Scale 2.54 ± 2.61; MDD, Inventory of Depressive Symptomatology Self-Report 46.67 ± 10.33.

**Supplementary Table 2.**
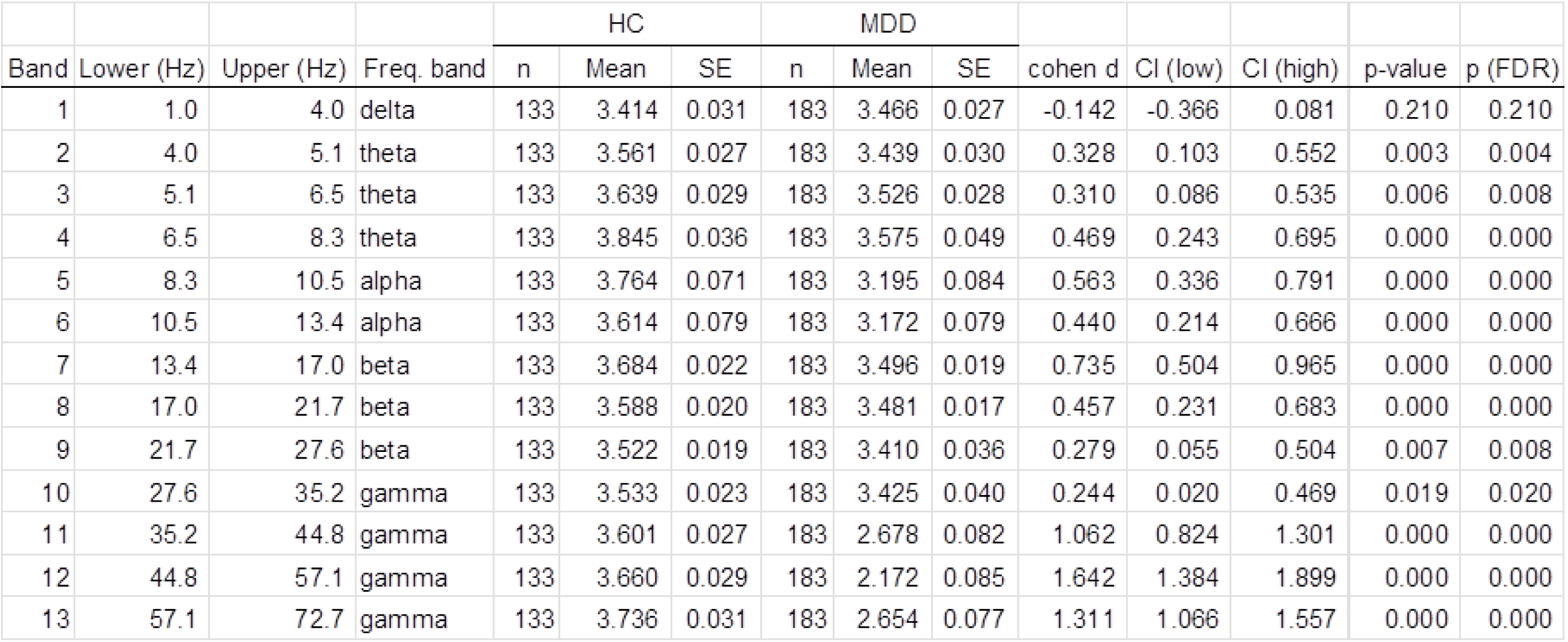
Unadjusted group comparison statistics: Bistability (BiS). Summary statistics from the unadjusted comparison between major depressive disorder (MDD) and healthy controls (HC). For each subject, metric values were averaged across channels. Group-level comparisons between MDD and HC were performed separately for each frequency band using Welch’s unequal-variance t-tests. Resulting p-values were adjusted for multiple testing across the 13 frequency bands within each metric using the Benjamini– Hochberg false discovery rate (FDR) procedure. Reported statistics include group means (±SE) for HC and MDD, effect sizes as Cohen’s d (MDD − HC), and corresponding 95% confidence intervals for d. n indicates the number of subjects included in each group for a given frequency band. Abbreviations: HC, healthy controls; MDD, major depressive disorder; SE, standard error; CI, confidence interval; FDR, false discovery rate; Freq. Band, frequency band.

**Supplementary Table 3.** Unadjusted group comparison statistics: Detrended fluctuation analysis (DFA). Statistical analysis identical to that described for Supplementary Table 2 (BiS). Reported statistics include group means (±SE) for HC and MDD, effect sizes as Cohen’s d (MDD − HC), 95% confidence intervals for d, and p- and FDR-adjusted p-values.

|  |  |  |  | HC |  |  | MDD |  |  |  |  |  |  |  |
| --- | --- | --- | --- | --- | --- | --- | --- | --- | --- | --- | --- | --- | --- | --- |
| Band | Lower (Hz) | Upper (Hz) | Freq. band | n | Mean | SE | n | Mean | SE | cohen d | CI (low) | CI (high) | p-value | p (FDR) |
| 1 | 1.0 | 4.0 | delta | 133 | 0.610 | 0.005 | 183 | 0.613 | 0.004 | -0.049 | -0.272 | 0.175 | 0.669 | 0.669 |
| 2 | 4.0 | 5.1 | theta | 133 | 0.602 | 0.005 | 183 | 0.599 | 0.004 | 0.050 | -0.174 | 0.273 | 0.659 | 0.669 |
| 3 | 5.1 | 6.5 | theta | 133 | 0.619 | 0.006 | 183 | 0.615 | 0.005 | 0.053 | -0.170 | 0.276 | 0.640 | 0.669 |
| 4 | 6.5 | 8.3 | theta | 133 | 0.686 | 0.009 | 183 | 0.659 | 0.007 | 0.283 | 0.058 | 0.507 | 0.014 | 0.019 |
| 5 | 8.3 | 10.5 | alpha | 133 | 0.741 | 0.008 | 183 | 0.680 | 0.007 | 0.625 | 0.396 | 0.853 | 0.000 | 0.000 |
| 6 | 10.5 | 13.4 | alpha | 133 | 0.739 | 0.008 | 183 | 0.676 | 0.007 | 0.693 | 0.464 | 0.923 | 0.000 | 0.000 |
| 7 | 13.4 | 17.0 | beta | 133 | 0.682 | 0.006 | 183 | 0.649 | 0.005 | 0.511 | 0.284 | 0.737 | 0.000 | 0.000 |
| 8 | 17.0 | 21.7 | beta | 133 | 0.676 | 0.005 | 183 | 0.638 | 0.004 | 0.657 | 0.428 | 0.886 | 0.000 | 0.000 |
| 9 | 21.7 | 27.6 | beta | 133 | 0.665 | 0.004 | 183 | 0.634 | 0.004 | 0.614 | 0.386 | 0.842 | 0.000 | 0.000 |
| 10 | 27.6 | 35.2 | gamma | 133 | 0.649 | 0.004 | 183 | 0.634 | 0.003 | 0.337 | 0.112 | 0.562 | 0.003 | 0.005 |
| 11 | 35.2 | 44.8 | gamma | 133 | 0.658 | 0.005 | 183 | 0.628 | 0.004 | 0.535 | 0.308 | 0.762 | 0.000 | 0.000 |
| 12 | 44.8 | 57.1 | gamma | 133 | 0.672 | 0.005 | 183 | 0.643 | 0.005 | 0.481 | 0.254 | 0.707 | 0.000 | 0.000 |
| 13 | 57.1 | 72.7 | gamma | 133 | 0.686 | 0.005 | 183 | 0.665 | 0.005 | 0.331 | 0.106 | 0.556 | 0.003 | 0.005 |

**Supplementary Table 4.** Unadjusted group comparison statistics: High-to-low power ratio (E/I_HLP_). Statistical analysis identical to that described for Supplementary Table 2 (BiS). Reported statistics include group means (±SE) for HC and MDD, effect sizes as Cohen’s d (MDD − HC), 95% confidence intervals for d, and p- and FDR-adjusted p-values.

|  |  |  |  | HC |  |  | MDD |  |  |  |  |  |  |  |
| --- | --- | --- | --- | --- | --- | --- | --- | --- | --- | --- | --- | --- | --- | --- |
| Band | Lower (Hz) | Upper (Hz) | Freq. band | n | Mean | SE | n | Mean | SE | cohen d | CI (low) | CI (high) | p-value | p (FDR) |
| 1 | 1.0 | 4.0 | delta | 133 | 0.254 | 0.008 | 183 | 0.261 | 0.007 | -0.068 | -0.292 | 0.155 | 0.548 | 0.647 |
| 2 | 4.0 | 5.1 | theta | 133 | 0.293 | 0.009 | 183 | 0.340 | 0.009 | -0.421 | -0.646 | -0.195 | 0.000 | 0.001 |
| 3 | 5.1 | 6.5 | theta | 133 | 0.299 | 0.008 | 183 | 0.358 | 0.009 | -0.526 | -0.753 | -0.299 | 0.000 | 0.000 |
| 4 | 6.5 | 8.3 | theta | 133 | 0.378 | 0.013 | 182 | 0.412 | 0.011 | -0.222 | -0.446 | 0.002 | 0.053 | 0.087 |
| 5 | 8.3 | 10.5 | alpha | 132 | 0.505 | 0.017 | 182 | 0.501 | 0.013 | 0.020 | -0.204 | 0.244 | 0.866 | 0.919 |
| 6 | 10.5 | 13.4 | alpha | 133 | 0.497 | 0.016 | 180 | 0.495 | 0.013 | 0.012 | -0.212 | 0.236 | 0.919 | 0.919 |
| 7 | 13.4 | 17.0 | beta | 133 | 0.428 | 0.012 | 183 | 0.460 | 0.009 | -0.248 | -0.472 | -0.024 | 0.032 | 0.059 |
| 8 | 17.0 | 21.7 | beta | 133 | 0.462 | 0.011 | 183 | 0.477 | 0.009 | -0.120 | -0.343 | 0.104 | 0.300 | 0.434 |
| 9 | 21.7 | 27.6 | beta | 133 | 0.430 | 0.009 | 180 | 0.418 | 0.010 | 0.095 | -0.129 | 0.319 | 0.390 | 0.508 |
| 10 | 27.6 | 35.2 | gamma | 133 | 0.337 | 0.009 | 180 | 0.308 | 0.009 | 0.254 | 0.029 | 0.479 | 0.024 | 0.052 |
| 11 | 35.2 | 44.8 | gamma | 133 | 0.288 | 0.009 | 177 | 0.201 | 0.009 | 0.765 | 0.532 | 0.998 | 0.000 | 0.000 |
| 12 | 44.8 | 57.1 | gamma | 133 | 0.264 | 0.010 | 177 | 0.158 | 0.008 | 0.995 | 0.757 | 1.233 | 0.000 | 0.000 |
| 13 | 57.1 | 72.7 | gamma | 133 | 0.252 | 0.010 | 181 | 0.182 | 0.008 | 0.634 | 0.404 | 0.863 | 0.000 | 0.000 |

**Supplementary Table 5.**
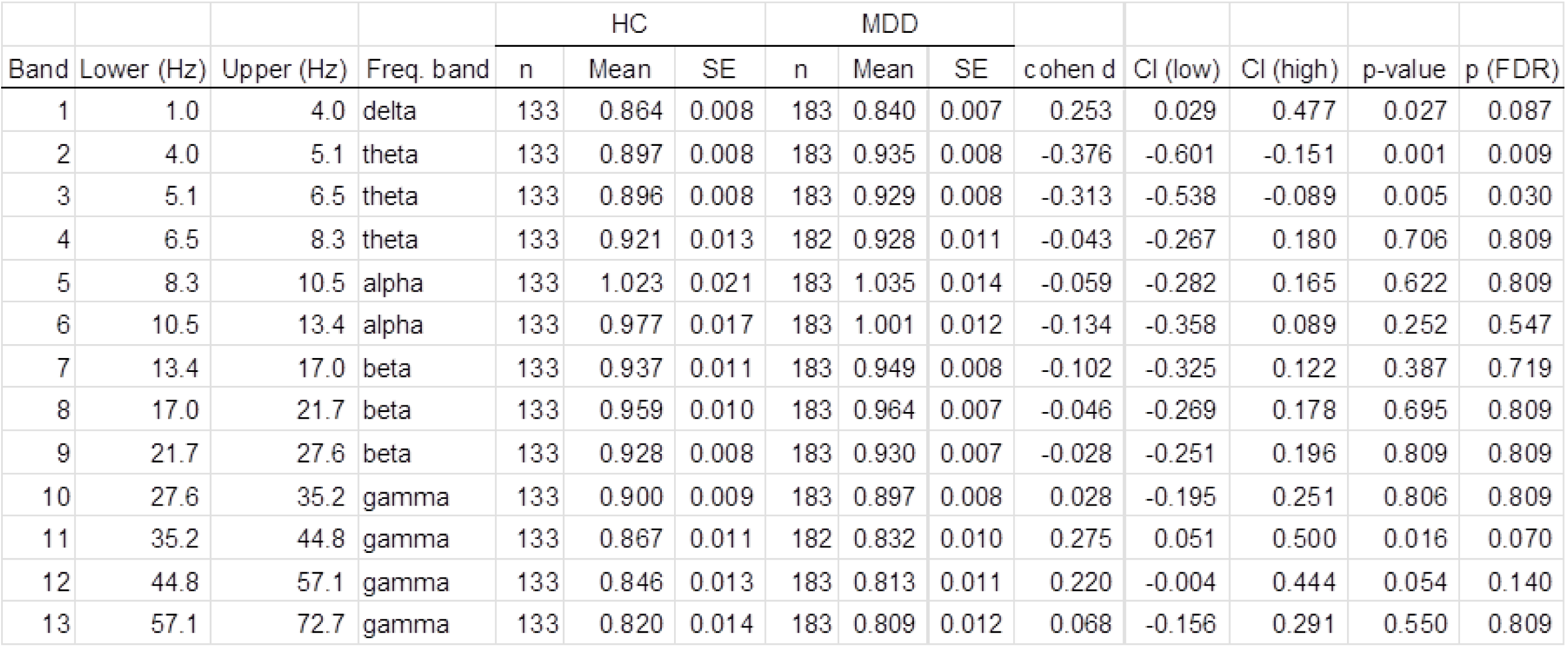
Unadjusted group comparison statistics: functional E/I ratio (fE/I). Statistical analysis identical to that described for Supplementary Table 2 (BiS). Reported statistics include group means (±SE) for HC and MDD, effect sizes as Cohen’s d (MDD − HC), 95% confidence intervals for d, and p- and FDR-adjusted p-values.

|  |  |  |  | HC |  |  | MDD |  |  |  |  |  |  |  |
| --- | --- | --- | --- | --- | --- | --- | --- | --- | --- | --- | --- | --- | --- | --- |
| Band | Lower (Hz) | Upper (Hz) | Freq. band | n | Mean | SE | n | Mean | SE | cohen d | CI (low) | CI (high) | p-value | p (FDR) |
| 1 | 1.0 | 4.0 | delta | 133 | 0.864 | 0.008 | 183 | 0.840 | 0.007 | 0.253 | 0.029 | 0.477 | 0.027 | 0.087 |
| 2 | 4.0 | 5.1 | theta | 133 | 0.897 | 0.008 | 183 | 0.935 | 0.008 | -0.376 | -0.601 | -0.151 | 0.001 | 0.009 |
| 3 | 5.1 | 6.5 | theta | 133 | 0.896 | 0.008 | 183 | 0.929 | 0.008 | -0.313 | -0.538 | -0.089 | 0.005 | 0.030 |
| 4 | 6.5 | 8.3 | theta | 133 | 0.921 | 0.013 | 182 | 0.928 | 0.011 | -0.043 | -0.267 | 0.180 | 0.706 | 0.809 |
| 5 | 8.3 | 10.5 | alpha | 133 | 1.023 | 0.021 | 183 | 1.035 | 0.014 | -0.059 | -0.282 | 0.165 | 0.622 | 0.809 |
| 6 | 10.5 | 13.4 | alpha | 133 | 0.977 | 0.017 | 183 | 1.001 | 0.012 | -0.134 | -0.358 | 0.089 | 0.252 | 0.547 |
| 7 | 13.4 | 17.0 | beta | 133 | 0.937 | 0.011 | 183 | 0.949 | 0.008 | -0.102 | -0.325 | 0.122 | 0.387 | 0.719 |
| 8 | 17.0 | 21.7 | beta | 133 | 0.959 | 0.010 | 183 | 0.964 | 0.007 | -0.046 | -0.269 | 0.178 | 0.695 | 0.809 |
| 9 | 21.7 | 27.6 | beta | 133 | 0.928 | 0.008 | 183 | 0.930 | 0.007 | -0.028 | -0.251 | 0.196 | 0.809 | 0.809 |
| 10 | 27.6 | 35.2 | gamma | 133 | 0.900 | 0.009 | 183 | 0.897 | 0.008 | 0.028 | -0.195 | 0.251 | 0.806 | 0.809 |
| 11 | 35.2 | 44.8 | gamma | 133 | 0.867 | 0.011 | 182 | 0.832 | 0.010 | 0.275 | 0.051 | 0.500 | 0.016 | 0.070 |
| 12 | 44.8 | 57.1 | gamma | 133 | 0.846 | 0.013 | 183 | 0.813 | 0.011 | 0.220 | -0.004 | 0.444 | 0.054 | 0.140 |
| 13 | 57.1 | 72.7 | gamma | 133 | 0.820 | 0.014 | 183 | 0.809 | 0.012 | 0.068 | -0.156 | 0.291 | 0.550 | 0.809 |

**Supplementary Table 6.** Unadjusted group comparison statistics: Separation of high- and low-power oscillations (E+I_HLS_). Statistical analysis identical to that described for Supplementary Table 2 (BiS). Reported statistics include group means (±SE) for HC and MDD, effect sizes as Cohen’s d (MDD − HC), 95% confidence intervals for d, and p- and FDR-adjusted p-values.

|  |  |  |  | HC |  |  | MDD |  |  |  |  |  |  |  |
| --- | --- | --- | --- | --- | --- | --- | --- | --- | --- | --- | --- | --- | --- | --- |
| Band | Lower (Hz) | Upper (Hz) | Freq. band | n | Mean | SE | n | Mean | SE | cohen d | CI (low) | CI (high) | p-value | p (FDR) |
| 1 | 1.0 | 4.0 | delta | 133 | 0.493 | 0.009 | 183 | 0.517 | 0.008 | -0.215 | -0.438 | 0.009 | 0.058 | 0.083 |
| 2 | 4.0 | 5.1 | theta | 133 | 0.498 | 0.008 | 183 | 0.480 | 0.008 | 0.165 | -0.059 | 0.388 | 0.138 | 0.180 |
| 3 | 5.1 | 6.5 | theta | 133 | 0.507 | 0.009 | 183 | 0.490 | 0.008 | 0.162 | -0.061 | 0.386 | 0.153 | 0.180 |
| 4 | 6.5 | 8.3 | theta | 133 | 0.609 | 0.014 | 182 | 0.548 | 0.010 | 0.410 | 0.184 | 0.636 | 0.000 | 0.001 |
| 5 | 8.3 | 10.5 | alpha | 132 | 0.760 | 0.020 | 182 | 0.606 | 0.017 | 0.657 | 0.427 | 0.887 | 0.000 | 0.000 |
| 6 | 10.5 | 13.4 | alpha | 133 | 0.666 | 0.017 | 180 | 0.542 | 0.014 | 0.638 | 0.408 | 0.867 | 0.000 | 0.000 |
| 7 | 13.4 | 17.0 | beta | 133 | 0.509 | 0.008 | 183 | 0.444 | 0.006 | 0.762 | 0.531 | 0.993 | 0.000 | 0.000 |
| 8 | 17.0 | 21.7 | beta | 133 | 0.479 | 0.006 | 183 | 0.424 | 0.006 | 0.729 | 0.499 | 0.960 | 0.000 | 0.000 |
| 9 | 21.7 | 27.6 | beta | 133 | 0.431 | 0.007 | 180 | 0.407 | 0.006 | 0.306 | 0.080 | 0.531 | 0.008 | 0.013 |
| 10 | 27.6 | 35.2 | gamma | 133 | 0.440 | 0.009 | 180 | 0.434 | 0.008 | 0.059 | -0.166 | 0.283 | 0.605 | 0.605 |
| 11 | 35.2 | 44.8 | gamma | 133 | 0.489 | 0.010 | 177 | 0.506 | 0.010 | -0.138 | -0.363 | 0.088 | 0.226 | 0.245 |
| 12 | 44.8 | 57.1 | gamma | 133 | 0.536 | 0.011 | 177 | 0.578 | 0.010 | -0.311 | -0.538 | -0.085 | 0.006 | 0.013 |
| 13 | 57.1 | 72.7 | gamma | 133 | 0.571 | 0.011 | 181 | 0.614 | 0.011 | -0.298 | -0.523 | -0.073 | 0.008 | 0.013 |

**Supplementary Table 7.** Covariate-adjusted group comparison statistics: Bistability (BiS). Summary statistics from the covariate-adjusted comparison between major depressive disorder (MDD) and healthy controls (HC). For each subject, metric values were averaged across channels. Group-level analyses were performed separately for each frequency band using robust linear regression models with group (MDD vs. HC) as the predictor of interest and age (z-scored) and sex (male = 1, female = 0) as covariates. To obtain covariate-adjusted group differences, group coefficients and their corresponding t-statistics were extracted from the full model. Reported statistics include covariate-adjusted group means (±SE) for HC and MDD, effect sizes as Cohen’s d (MDD − HC), and corresponding 95% confidence intervals for d. n indicates the number of subjects included in each group for a given frequency band. Resulting p-values were adjusted for multiple testing across the 13 frequency bands within each metric using the Benjamini–Hochberg false discovery rate (FDR) procedure. Abbreviations: HC, healthy controls; MDD, major depressive disorder; SE, standard error; CI, confidence interval; FDR, false discovery rate; Freq. Band, frequency band.

|  |  |  |  | HC |  |  | MDD |  |  |  |  |  |  |  |
| --- | --- | --- | --- | --- | --- | --- | --- | --- | --- | --- | --- | --- | --- | --- |
| Band | Lower (Hz) | Upper (Hz) | Freq. band | n | Mean | SE | n | Mean | SE | cohen d | CI (low) | CI (high) | p-value | p (FDR) |
| 1 | 1.0 | 4.0 | delta | 133 | -0.023 | 0.026 | 183 | 0.017 | 0.022 | -0.077 | 0.369 | 0.145 | 0.201 | 0.238 |
| 2 | 4.0 | 5.1 | theta | 133 | 0.052 | 0.022 | 183 | -0.039 | 0.021 | -0.603 | -0.150 | -0.374 | 0.001 | 0.002 |
| 3 | 5.1 | 6.5 | theta | 133 | 0.049 | 0.026 | 183 | -0.035 | 0.022 | -0.528 | -0.078 | -0.301 | 0.008 | 0.011 |
| 4 | 6.5 | 8.3 | theta | 133 | 0.082 | 0.032 | 183 | -0.063 | 0.028 | -0.646 | -0.191 | -0.416 | 0.000 | 0.000 |
| 5 | 8.3 | 10.5 | alpha | 133 | 0.134 | 0.045 | 183 | -0.111 | 0.041 | -0.727 | -0.268 | -0.495 | 0.000 | 0.000 |
| 6 | 10.5 | 13.4 | alpha | 133 | 0.134 | 0.039 | 183 | -0.105 | 0.032 | -0.831 | -0.366 | -0.595 | 0.000 | 0.000 |
| 7 | 13.4 | 17.0 | beta | 133 | 0.088 | 0.019 | 183 | -0.063 | 0.016 | -1.003 | -0.527 | -0.760 | 0.000 | 0.000 |
| 8 | 17.0 | 21.7 | beta | 133 | 0.038 | 0.016 | 183 | -0.026 | 0.012 | -0.647 | -0.192 | -0.417 | 0.000 | 0.000 |
| 9 | 21.7 | 27.6 | beta | 133 | 0.005 | 0.016 | 183 | -0.004 | 0.013 | -0.281 | 0.165 | -0.057 | 0.613 | 0.664 |
| 10 | 27.6 | 35.2 | gamma | 133 | 0.005 | 0.017 | 183 | -0.004 | 0.015 | -0.265 | 0.181 | -0.042 | 0.712 | 0.712 |
| 11 | 35.2 | 44.8 | gamma | 133 | 0.118 | 0.024 | 183 | -0.127 | 0.039 | -0.870 | -0.403 | -0.632 | 0.000 | 0.000 |
| 12 | 44.8 | 57.1 | gamma | 133 | 0.767 | 0.032 | 183 | -0.591 | 0.082 | -1.999 | -1.415 | -1.696 | 0.000 | 0.000 |
| 13 | 57.1 | 72.7 | gamma | 133 | 0.516 | 0.031 | 183 | -0.411 | 0.072 | -1.501 | -0.978 | -1.232 | 0.000 | 0.000 |

**Supplementary Table 8.** Covariate-adjusted group comparison statistics: Detrended fluctuation analysis (DFA). Statistical analysis was identical to that described for Supplementary Table 7 (BiS). Reported statistics include group means (±SE) for HC and MDD, effect sizes as Cohen’s *d* (MDD − HC), 95% confidence intervals for *d*, and *p*- and FDR-adjusted *p*-values.

|  |  |  |  | HC |  |  | MDD |  |  |  |  |  |  |  |
| --- | --- | --- | --- | --- | --- | --- | --- | --- | --- | --- | --- | --- | --- | --- |
| Band | Lower (Hz) | Upper (Hz) | Freq. band | n | Mean | SE | n | Mean | SE | cohen d | CI (low) | CI (high) | p-value | p (FDR) |
| 1 | 1.0 | 4.0 | delta | 133 | 0.007 | 0.005 | 183 | 0.010 | 0.004 | 0.045 | -0.009 | 0.014 | 0.417 | 0.493 |
| 2 | 4.0 | 5.1 | theta | 133 | 0.009 | 0.005 | 183 | 0.006 | 0.004 | -0.057 | -0.016 | 0.010 | 0.589 | 0.589 |
| 3 | 5.1 | 6.5 | theta | 133 | 0.009 | 0.006 | 183 | 0.006 | 0.005 | -0.052 | -0.019 | 0.012 | 0.554 | 0.589 |
| 4 | 6.5 | 8.3 | theta | 133 | 0.024 | 0.008 | 183 | -0.002 | 0.007 | -0.270 | -0.047 | -0.004 | 0.014 | 0.019 |
| 5 | 8.3 | 10.5 | alpha | 133 | 0.044 | 0.008 | 183 | -0.017 | 0.007 | -0.617 | -0.082 | -0.039 | 0.000 | 0.001 |
| 6 | 10.5 | 13.4 | alpha | 133 | 0.044 | 0.008 | 183 | -0.017 | 0.007 | -0.679 | -0.081 | -0.041 | 0.000 | 0.001 |
| 7 | 13.4 | 17.0 | beta | 133 | 0.028 | 0.006 | 183 | -0.006 | 0.005 | -0.515 | -0.047 | -0.019 | 0.000 | 0.001 |
| 8 | 17.0 | 21.7 | beta | 133 | 0.031 | 0.005 | 183 | -0.008 | 0.004 | -0.658 | -0.052 | -0.026 | 0.000 | 0.001 |
| 9 | 21.7 | 27.6 | beta | 133 | 0.024 | 0.004 | 183 | -0.006 | 0.004 | -0.612 | -0.041 | -0.019 | 0.000 | 0.001 |
| 10 | 27.6 | 35.2 | gamma | 133 | 0.015 | 0.004 | 183 | -0.001 | 0.003 | -0.340 | -0.026 | -0.005 | 0.003 | 0.004 |
| 11 | 35.2 | 44.8 | gamma | 133 | 0.023 | 0.005 | 183 | -0.006 | 0.004 | -0.528 | -0.041 | -0.017 | 0.000 | 0.001 |
| 12 | 44.8 | 57.1 | gamma | 133 | 0.025 | 0.005 | 183 | -0.003 | 0.005 | -0.472 | -0.042 | -0.015 | 0.000 | 0.001 |
| 13 | 57.1 | 72.7 | gamma | 133 | 0.021 | 0.005 | 183 | 0.000 | 0.005 | -0.323 | -0.035 | -0.007 | 0.000 | 0.001 |

**Supplementary Table 9.** Covariate-adjusted group comparison statistics: High-to-low power ratio (E/I_HLP_). Statistical analysis was identical to that described for Supplementary Table 7 (BiS). Reported statistics include group means (±SE) for HC and MDD, effect sizes as Cohen’s *d* (MDD − HC), 95% confidence intervals for *d*, and *p*- and FDR-adjusted *p*-values.

|  |  |  |  | HC |  |  | MDD |  |  |  |  |  |  |  |
| --- | --- | --- | --- | --- | --- | --- | --- | --- | --- | --- | --- | --- | --- | --- |
| Band | Lower (Hz) | Upper (Hz) | Freq. band | n | Mean | SE | n | Mean | SE | cohen d | CI (low) | CI (high) | p-value | p (FDR) |
| 1 | 1.0 | 4.0 | delta | 133 | -0.003 | 0.007 | 183 | 0.002 | 0.006 | -0.162 | 0.284 | 0.061 | 0.591 | 0.698 |
| 2 | 4.0 | 5.1 | theta | 133 | -0.025 | 0.008 | 183 | 0.019 | 0.008 | 0.214 | 0.670 | 0.440 | 0.000 | 0.000 |
| 3 | 5.1 | 6.5 | theta | 133 | -0.029 | 0.008 | 183 | 0.022 | 0.008 | 0.299 | 0.759 | 0.526 | 0.000 | 0.000 |
| 4 | 6.5 | 8.3 | theta | 133 | -0.020 | 0.012 | 182 | 0.014 | 0.010 | 0.033 | 0.483 | 0.257 | 0.024 | 0.045 |
| 5 | 8.3 | 10.5 | alpha | 132 | -0.001 | 0.016 | 182 | 0.001 | 0.013 | -0.214 | 0.233 | 0.009 | 0.935 | 0.950 |
| 6 | 10.5 | 13.4 | alpha | 133 | -0.001 | 0.015 | 180 | 0.000 | 0.012 | -0.217 | 0.231 | 0.007 | 0.950 | 0.950 |
| 7 | 13.4 | 17.0 | beta | 133 | -0.015 | 0.010 | 183 | 0.011 | 0.009 | 0.020 | 0.468 | 0.243 | 0.033 | 0.054 |
| 8 | 17.0 | 21.7 | beta | 133 | -0.006 | 0.010 | 183 | 0.004 | 0.008 | -0.127 | 0.319 | 0.095 | 0.401 | 0.521 |
| 9 | 21.7 | 27.6 | beta | 133 | 0.007 | 0.008 | 180 | -0.006 | 0.009 | -0.353 | 0.095 | -0.128 | 0.261 | 0.377 |
| 10 | 27.6 | 35.2 | gamma | 133 | 0.016 | 0.009 | 180 | -0.012 | 0.008 | -0.511 | -0.059 | -0.284 | 0.013 | 0.029 |
| 11 | 35.2 | 44.8 | gamma | 133 | 0.048 | 0.008 | 177 | -0.036 | 0.008 | -1.162 | -0.668 | -0.909 | 0.000 | 0.000 |
| 12 | 44.8 | 57.1 | gamma | 133 | 0.054 | 0.008 | 177 | -0.039 | 0.007 | -1.341 | -0.830 | -1.079 | 0.000 | 0.000 |
| 13 | 57.1 | 72.7 | gamma | 133 | 0.037 | 0.009 | 181 | -0.026 | 0.007 | -0.924 | -0.451 | -0.683 | 0.000 | 0.000 |

**Supplementary Table 10.** Covariate-adjusted group comparison statistics: functional E/I ratio (fE/I). Statistical analysis was identical to that described for Supplementary Table 7 (BiS). Reported statistics include group means (±SE) for HC and MDD, effect sizes as Cohen’s *d* (MDD − HC), 95% confidence intervals for *d*, and *p*- and FDR-adjusted *p*-values.

|  |  |  |  | HC |  |  | MDD |  |  |  |  |  |  |  |
| --- | --- | --- | --- | --- | --- | --- | --- | --- | --- | --- | --- | --- | --- | --- |
| Band | Lower (Hz) | Upper (Hz) | Freq. band | n | Mean | SE | n | Mean | SE | cohen d | CI (low) | CI (high) | p-value | p (FDR) |
| 1 | 1.0 | 4.0 | delta | 133 | 0.010 | 0.007 | 183 | -0.007 | 0.006 | -0.449 | -0.001 | -0.223 | 0.049 | 0.128 |
| 2 | 4.0 | 5.1 | theta | 133 | -0.025 | 0.007 | 183 | 0.019 | 0.006 | 0.394 | 0.861 | 0.624 | 0.000 | 0.000 |
| 3 | 5.1 | 6.5 | theta | 133 | -0.020 | 0.007 | 183 | 0.015 | 0.007 | 0.240 | 0.697 | 0.465 | 0.000 | 0.000 |
| 4 | 6.5 | 8.3 | theta | 133 | -0.006 | 0.011 | 182 | 0.004 | 0.009 | -0.133 | 0.313 | 0.089 | 0.431 | 0.497 |
| 5 | 8.3 | 10.5 | alpha | 133 | -0.010 | 0.018 | 183 | 0.007 | 0.012 | -0.122 | 0.324 | 0.101 | 0.375 | 0.497 |
| 6 | 10.5 | 13.4 | alpha | 133 | -0.014 | 0.014 | 183 | 0.010 | 0.011 | -0.046 | 0.401 | 0.176 | 0.120 | 0.261 |
| 7 | 13.4 | 17.0 | beta | 133 | -0.005 | 0.010 | 183 | 0.004 | 0.007 | -0.118 | 0.328 | 0.104 | 0.359 | 0.497 |
| 8 | 17.0 | 21.7 | beta | 133 | -0.001 | 0.009 | 183 | 0.000 | 0.006 | -0.212 | 0.234 | 0.011 | 0.923 | 0.923 |
| 9 | 21.7 | 27.6 | beta | 133 | -0.004 | 0.007 | 183 | 0.003 | 0.006 | -0.125 | 0.321 | 0.098 | 0.390 | 0.497 |
| 10 | 27.6 | 35.2 | gamma | 133 | -0.004 | 0.007 | 183 | 0.003 | 0.006 | -0.138 | 0.307 | 0.084 | 0.459 | 0.497 |
| 11 | 35.2 | 44.8 | gamma | 133 | 0.019 | 0.009 | 182 | -0.014 | 0.008 | -0.550 | -0.098 | -0.322 | 0.005 | 0.021 |
| 12 | 44.8 | 57.1 | gamma | 133 | 0.019 | 0.011 | 183 | -0.014 | 0.010 | -0.493 | -0.043 | -0.266 | 0.019 | 0.062 |
| 13 | 57.1 | 72.7 | gamma | 133 | 0.008 | 0.013 | 183 | -0.006 | 0.011 | -0.328 | 0.118 | -0.104 | 0.358 | 0.497 |

**Supplementary Table 11.**
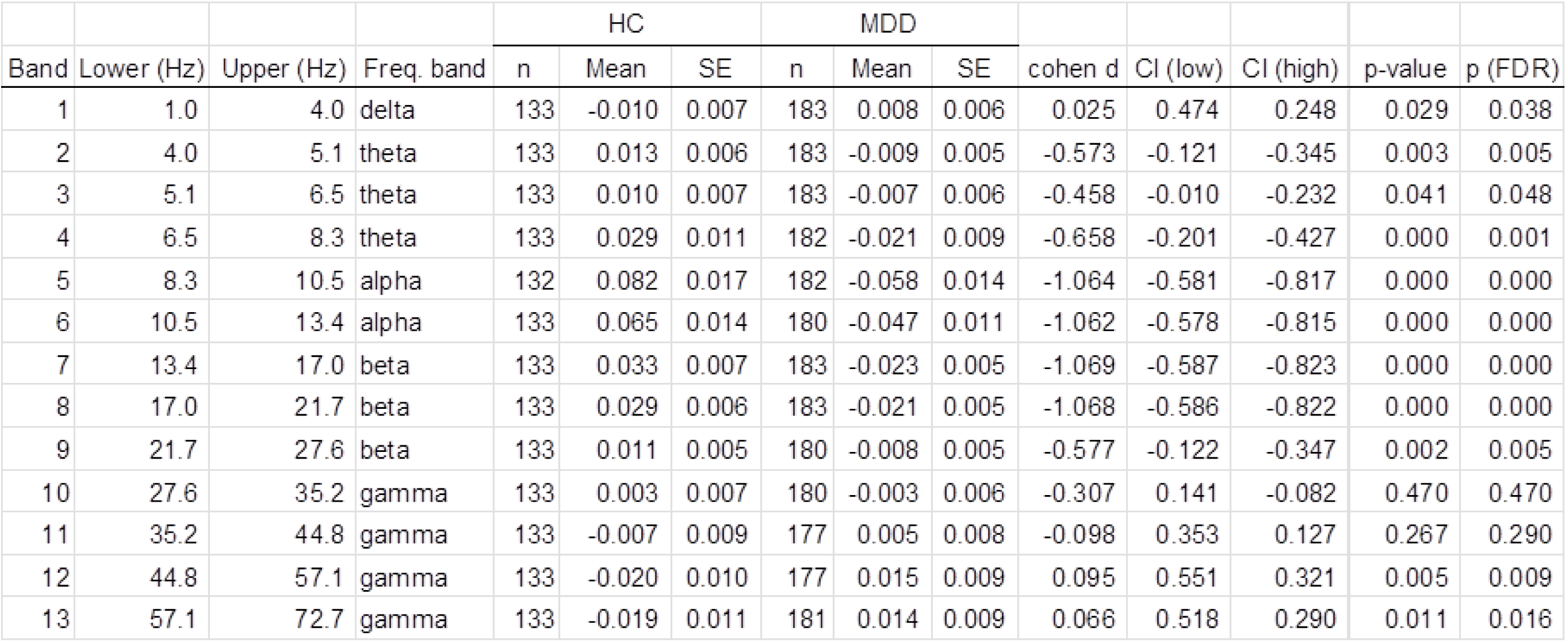
Covariate-adjusted group comparison statistics: Separation of high- and low-power oscillations (E+I_HLS_). Statistical analysis was identical to that described for Supplementary Table 7 (BiS). Reported statistics include group means (±SE) for HC and MDD, effect sizes as Cohen’s *d* (MDD − HC), 95% confidence intervals for *d*, and *p*- and FDR-adjusted *p*-values.

|  |  |  |  | HC |  |  | MDD |  |  |  |  |  |  |  |
| --- | --- | --- | --- | --- | --- | --- | --- | --- | --- | --- | --- | --- | --- | --- |
| Band | Lower (Hz) | Upper (Hz) | Freq. band | n | Mean | SE | n | Mean | SE | cohen d | CI (low) | CI (high) | p-value | p (FDR) |
| 1 | 1.0 | 4.0 | delta | 133 | -0.010 | 0.007 | 183 | 0.008 | 0.006 | 0.025 | 0.474 | 0.248 | 0.029 | 0.038 |
| 2 | 4.0 | 5.1 | theta | 133 | 0.013 | 0.006 | 183 | -0.009 | 0.005 | -0.573 | -0.121 | -0.345 | 0.003 | 0.005 |
| 3 | 5.1 | 6.5 | theta | 133 | 0.010 | 0.007 | 183 | -0.007 | 0.006 | -0.458 | -0.010 | -0.232 | 0.041 | 0.048 |
| 4 | 6.5 | 8.3 | theta | 133 | 0.029 | 0.011 | 182 | -0.021 | 0.009 | -0.658 | -0.201 | -0.427 | 0.000 | 0.001 |
| 5 | 8.3 | 10.5 | alpha | 132 | 0.082 | 0.017 | 182 | -0.058 | 0.014 | -1.064 | -0.581 | -0.817 | 0.000 | 0.000 |
| 6 | 10.5 | 13.4 | alpha | 133 | 0.065 | 0.014 | 180 | -0.047 | 0.011 | -1.062 | -0.578 | -0.815 | 0.000 | 0.000 |
| 7 | 13.4 | 17.0 | beta | 133 | 0.033 | 0.007 | 183 | -0.023 | 0.005 | -1.069 | -0.587 | -0.823 | 0.000 | 0.000 |
| 8 | 17.0 | 21.7 | beta | 133 | 0.029 | 0.006 | 183 | -0.021 | 0.005 | -1.068 | -0.586 | -0.822 | 0.000 | 0.000 |
| 9 | 21.7 | 27.6 | beta | 133 | 0.011 | 0.005 | 180 | -0.008 | 0.005 | -0.577 | -0.122 | -0.347 | 0.002 | 0.005 |
| 10 | 27.6 | 35.2 | gamma | 133 | 0.003 | 0.007 | 180 | -0.003 | 0.006 | -0.307 | 0.141 | -0.082 | 0.470 | 0.470 |
| 11 | 35.2 | 44.8 | gamma | 133 | -0.007 | 0.009 | 177 | 0.005 | 0.008 | -0.098 | 0.353 | 0.127 | 0.267 | 0.290 |
| 12 | 44.8 | 57.1 | gamma | 133 | -0.020 | 0.010 | 177 | 0.015 | 0.009 | 0.095 | 0.551 | 0.321 | 0.005 | 0.009 |
| 13 | 57.1 | 72.7 | gamma | 133 | -0.019 | 0.011 | 181 | 0.014 | 0.009 | 0.066 | 0.518 | 0.290 | 0.011 | 0.016 |

